# Methylotrophy in the Mire: direct and indirect routes for methane production in thawing permafrost

**DOI:** 10.1101/2023.07.09.548291

**Authors:** Jared B. Ellenbogen, Mikayla A. Borton, Bridget B. McGivern, Dylan R. Cronin, David W. Hoyt, Viviana Freire-Zapata, Carmody K. McCalley, Ruth K. Varner, Patrick M. Crill, Richard A. Wehr, Jeffrey P. Chanton, Ben J. Woodcroft, Malak M. Tfaily, Gene W. Tyson, Virginia I. Rich, Kelly C. Wrighton

## Abstract

While wetlands are major sources of biogenic methane (CH_4_), our understanding of resident microbial metabolisms is incomplete, which compromises prediction of CH_4_ emissions under ongoing climate change. Here, we employed genome-resolved multi-omics to expand our understanding of methanogenesis in the thawing permafrost peatland of Stordalen Mire, in arctic Sweden. In quadrupling the genomic representation of the site’s methanogens and examining their encoded metabolisms, we revealed that nearly 20% (72) of the metagenome-assembled genomes (MAGs) encoded potential for methylotrophic methanogenesis. Further, 27% of the transcriptionally active methanogens expressed methylotrophic genes; for *Methanosarcinales* and *Methanobacteriales* MAGs, these data indicated use of methylated oxygen compounds (e.g., methanol), while for *Methanomassiliicoccales*, they primarily implicated methyl sulfides and methylamines. In addition to methanogenic methylotrophy, >1700 bacterial MAGs across 19 phyla encoded anaerobic methylotrophic potential, with expression across 12 phyla. Metabolomic analyses revealed the presence of diverse methylated compounds in the Mire, including some known methylotrophic substrates. Active methylotrophy was observed across all stages of a permafrost thaw gradient in Stordalen, with the most frozen non-methanogenic palsa found to host bacterial methylotrophy, and the partially thawed bog and fully thawed fen seen to house both methanogenic and bacterial methylotrophic activity. Methanogenesis across increasing permafrost thaw is thus revised from sole dominance of hydrogenotrophic production, and the appearance of acetoclastic at full thaw, to consider co-occurrence of methylotrophy throughout. Collectively, these findings indicate that methanogenic and bacterial methylotrophy may be an important and previously underappreciated component of carbon cycling and emissions in these rapidly changing wetland habitats.

## Importance

Wetlands are the biggest natural source of atmospheric CH_4_ emissions, yet we have an incomplete understanding of the suite of microbial metabolisms that result in CH_4_ formation. Specifically, methanogenesis from methylated compounds is excluded from all ecosystem models used to predict wetland contributions to the global CH_4_ budget. Though recent studies have shown methylotrophic methanogenesis to be active across wetlands, the broad climatic importance of the metabolism remains critically understudied. Further, some methylotrophic bacteria are known to produce methanogenic by-products like acetate, increasing the complexity of the microbial methylotrophic metabolic network. Prior studies of Stordalen Mire have suggested methylotrophic methanogenesis irrelevant *in situ,* and not emphasized bacterial capacity for the metabolism, both of which we countered in this study. The importance of our findings lies in the significant advancement towards unraveling the broader impact of methylotrophs in wetland methanogenesis, and consequently, their contribution to the terrestrial global carbon cycle.

## Background

Wetlands are the largest natural source of atmospheric methane (CH_4_) emissions (1, 2), with those found in permafrost zones of specific concern to the global CH_4_ budget due to their sensitivity to a warming climate. Climate change-induced permafrost thaw is anticipated to make large quantities of previously frozen near-surface carbon available to soil microbiota over the next century, which could accelerate CH_4_ production and release (3). Methanogenic archaea are the primary biological producers of CH_4_ in soils, via three distinct pathways of methanogenesis (4). In saturated soils there is wide appreciation for hydrogenotrophic and acetoclastic pathways from microorganisms that utilize H_2_/carbon dioxide (CO_2_) and acetate, respectively, to form CH_4_ (4, 5). The methylotrophic pathway, which allows organisms to use methylated compounds for CH_4_ production (6, 7), is far less studied though appreciation of it is growing. This knowledge gap contributes to the fact that contemporary process-based biogeochemical models account only for hydrogenotrophic and acetoclastic microbial CH_4_ production in climate-relevant soil systems (8).

In contrast to historical paradigms, recent genomic insights have greatly expanded our understanding of the distribution and activity of methylotrophic methanogens, especially in saturated, high-CH_4_ emitting soils (9–17). Similarly, biochemical and physiological efforts have expanded the suite of substrates known to be utilized by methylotrophic methanogens (18–24). These methanogens catabolize methylated compounds via oxygen-sensitive substrate-specific so-called three-component methyltransferase systems (also frequently referred to as corrinoid-dependent methyltransferase systems) (6, 7). This gene content information can be used to physiologically classify methylotrophic methanogens as using one or more of three major substrate categories: methylated amines like trimethylamine or glycine betaine (methyl-N) (20, 25–27), methylated sulfides like dimethyl sulfide (methyl-S) (23, 28), and methylated oxygen compounds like methanol (methyl-O) (18, 19, 29). Moreover, methylotrophic methanogens can be obligate, meaning they only utilize methylated compounds to produce CH_4_, or facultative, meaning they encode and can express multiple pathways for CH_4_ production (10, 24). In addition to methanogens, some anaerobic bacteria can use similar corrinoid-dependent methyltransferase systems in a non-methanogenic mechanism for both carbon assimilation and energy generation (30–38). In summary, while diverse efforts have increased knowledge of anaerobic methylotrophic metabolisms, these metabolisms remain under characterized in climate relevant soil systems.

In this study, we used a genome-resolved, multi-omic approach to profile the potential for anaerobic, corrinoid-dependent methylotrophic metabolisms across a rapidly thawing CH_4_-emitting permafrost peatland (39, 40) located in arctic Sweden. This offered us a unique opportunity to study this metabolism across a discontinuous permafrost thaw gradient encompassing three distinct habitat types – palsa, bog, and fen – at multiple depths within each (Figure S1). Prior characterization of the soil microbiota across this thaw gradient has included analysis of the native methanogen community, focusing primarily on the dominant hydrogenotrophic *Methanoflorens stordalenmirensis* (genus “Bog-38” in the Genome Taxonomy Database (GTDB)) (41–43). Here, with metagenomic sequencing of more samples, we expand the catalog of methanogens for the site, offering new opportunities for resolved physiological characterization of methanogenic pathways. Using a combination of metagenome, metabolite, and metatranscriptome data we demonstrate anaerobic methylotrophy to be an underappreciated part of the CH_4_ cycle in Stordalen Mire and suggest the *in situ* methylotrophic metabolic network to be of previously unrecognized complexity. These findings let us build a new conceptual model of how methylotrophic metabolisms can directly and indirectly modulate CH_4_ fluxes across the Mire.

## Results

### One fifth of Stordalen Mire’s diverse methanogens, spanning 3 orders, encode methylotrophic methanogenesis

Field sampling from 2010-2017 has yielded an extensive microbial metagenome-assembled genome (MAG) database from Stordalen Mire (43). Of the MAGs with GTDB-assigned taxonomy of Archaea, functional analyses confirmed 367 (Table S1) encoded the potential for methanogenesis (Table S2). This roughly quadrupled the known methanogen MAGs for this site, with 23% of the methanogens having been previously reported in Woodcroft and Singleton et al. (43).

We next curated the metabolic potential of these MAGs, revealing a diversity of substrate-specific methanogenic potential (Figure S2, Table S2). To assess methylotrophic potential, we manually inspected MAGs for genes encoding three-component (or corrinoid-dependent) methyltransferase systems (Figure S3, Table S3), each comprised of a substrate:corrinoid methyltransferase (MtxB), a corrinoid-binding protein (MtxC), and a methylcorrinoid:carbon-carrier methyltransferase (MtxA), which together bring substrate-derived methyl groups into methanogenesis, and a reductive activase (RamX), which reactivates the corrinoid (Figure 1A). MAGs were analyzed considering both methyltransferase system component completeness and gene synteny, as *mtxBCA*/*ramX* genes are frequently co-encoded (30, 38, 44). This led us to identify 85 MAGs encoding a total of 438 methyltransferase system genes (Figure 1B, Table S2). These methyltransferase system genes were encoded in multiple representatives within the methanogenic archaeal orders, including all 5 MAGs within the *Methanomassiliicoccales*, all 70 MAGs within the *Methanobacteriales*, and within 10 (of 11) MAGs in the *Methanosarcinales*. In addition, we required a substrate-specific *mtxB* gene in all methanogenic MAGs considered methylotrophic, as we considered this to be the best single marker gene for the physiology, due to its encoding the enzyme that directly catalyzes substrate demethylation. This more conservative requirement retained all the *Methanomassiliicoccales*, 62 of the *Methanobacteriales,* and 5 of the *Methanosarcinales*. From genome potential, we reported that the *Methanosarcinales* and *Methanobacteriales* are likely facultative methylotrophs, and the *Methanomassiliicoccales* are likely obligate methylotrophs (Table S2). In summary we conservatively identified 72 methanogen MAGs with the potential to catalyze methylotrophy (Figure 1B).

**Figure 1.**
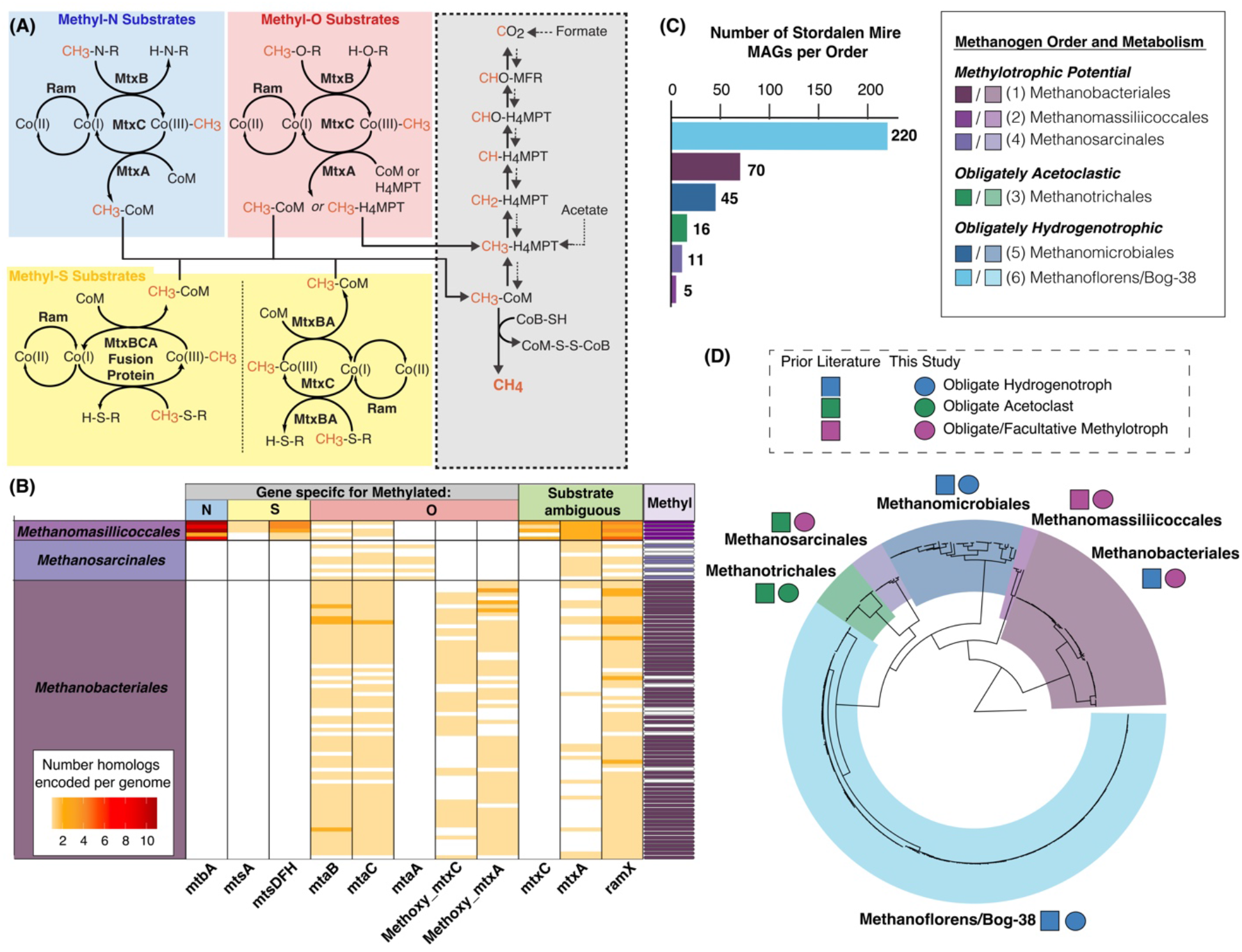
Phylogenomic and metagenomic analysis of Stordalen Mire methanogens. (A) Overview of substrate-specific physiology of methylotrophy, involving three-component (or corrinoid-dependent) methyltransferase systems to shuttle substrate-derived methyl groups into central methanogenesis (shown in grey). This includes a substrate:corrinoid methyltransferase (MtxB), a corrinoid-binding protein (MtxC), a methylated-corrinoid:carbon carrier methyltransferase (MtxA) and an activating enzyme (Ram or RamX). The internal “x” in the MtxABC protein/gene name (and the terminal “X” in RamX) is a generalized placeholder; the actual letter at this position varies to denote substrate specificity. Methyltransferase systems for methylated amines and oxygen compounds share a conserved architecture, while those for methylated sulfur compounds occur in two variations involving multi-functional proteins. (B) Heatmap showing the number of identified methylotrophic genes encoded in putative methylotrophic methanogen metagenome-assembled genomes (MAGs) from three orders. Each row represents a distinct MAG (grouped by taxonomic order), each column a different methylotrophic gene type, and cell color intensity denotes the number of identified genes. Gene columns are grouped by inferred substrate categories: methyl-N, -O, and -S, as in (A), and “substrate ambiguous” for those with annotation uncertainty or known substrate flexibility. Genes only identified to overall substrate category retain ’x’ in their names, and in the methyl-O substrates category, methoxylated substrates are indicated by the “methoxy” prefix. For more confident identifications of methylotrophic gene homologs, columns are named as substrate-specific methyltransferase system member genes (e.g., the methanol-specific *mtaB*). The furthest right column indicates whether each MAG meet the threshold criteria to be defined as methylotrophic (purple cell), or not (white cell). (C) Bar chart showing the number of Stordalen Mire derived methanogen MAGs per order present in data set (D) Overlay of phylogeny and genomically-inferred function for these 367 methanogen MAGs. MAGs were placed onto the GTDB r207 tree using 53 concatenated archaeal marker genes, and the tree was rooted with a GTDB-derived MAG from the archaeal phylum *Undinarchaeota*. Methanogen orders are delineated by color shading of the tree, and adjacent to each, genome-inferred methanogen pathway for the representatives at Stordalen Mire is denoted by colored squares for past metabolic designation, and circles for this study’s updated designation).

The *Methanomassiliicoccales* MAGs were found to encode genes for diverse methylotrophic substrates including methylated amines, methylated sulfides, and methanol (Figure 1B, Table S2). The *Methanobacteriales* and the *Methanosarcinales* encoded genes solely for demethylation of methylated oxygen compounds (e.g., methoxylated compounds or methanol), especially homologs of the methanol methyltransferase system components (Figure 1B). Though these latter two orders were previously considered hydrogenotrophic and acetoclastic, respectively, within Stordalen Mire (41, 42), they each include methylotrophic isolates providing support for our genomic inferences (24, 45–47). Our metabolic curation of the remainder of the methanogenic orders present in the site (Figure 1C-1D) was in agreement with prior metabolic assignments (41, 42). In total, these data refute the notion that the *Methanomassiliicoccales* are the only lineage within the Stordalen Mire methanogen community to encode potential for methylotrophic methanogenesis, highlighting the underappreciated potential of this metabolism.

### Methanogen relative abundance and diversity increases along the permafrost thaw gradient

The Mire contains three distinct habitat types which constitute a discontinuous, natural permafrost thaw gradient. In July 2016, we sampled the active layer in 3 habitat types: (i) palsa, overlaying intact permafrost, (ii) bog, with partially thawed permafrost and a perched water table, and (iii) fen, fully thawed and inundated (Figure S1, methods). CH_4_ flux increased with permafrost thaw state, from negligible from the palsa to highest from the fen (Figure 2A, Table S4). Consistent with this flux pattern, we failed to recruit reads to our methanogen MAGs from palsa metagenomes (Table S5), while the diversity of the methanogen orders (Figure 2B) and the total relative abundance of methanogens (Figure 2C) increased from bog to fen. Additionally, methanogen relative abundance increased significantly with depth in the bog, mirroring the water table depth, and likely reflecting saturated anoxic soil conditions favorable for methanogenesis (Figure 2B). Taken together, our findings support the idea that methanogens in Stordalen Mire chiefly reside in saturated soils.

**Figure 2.**
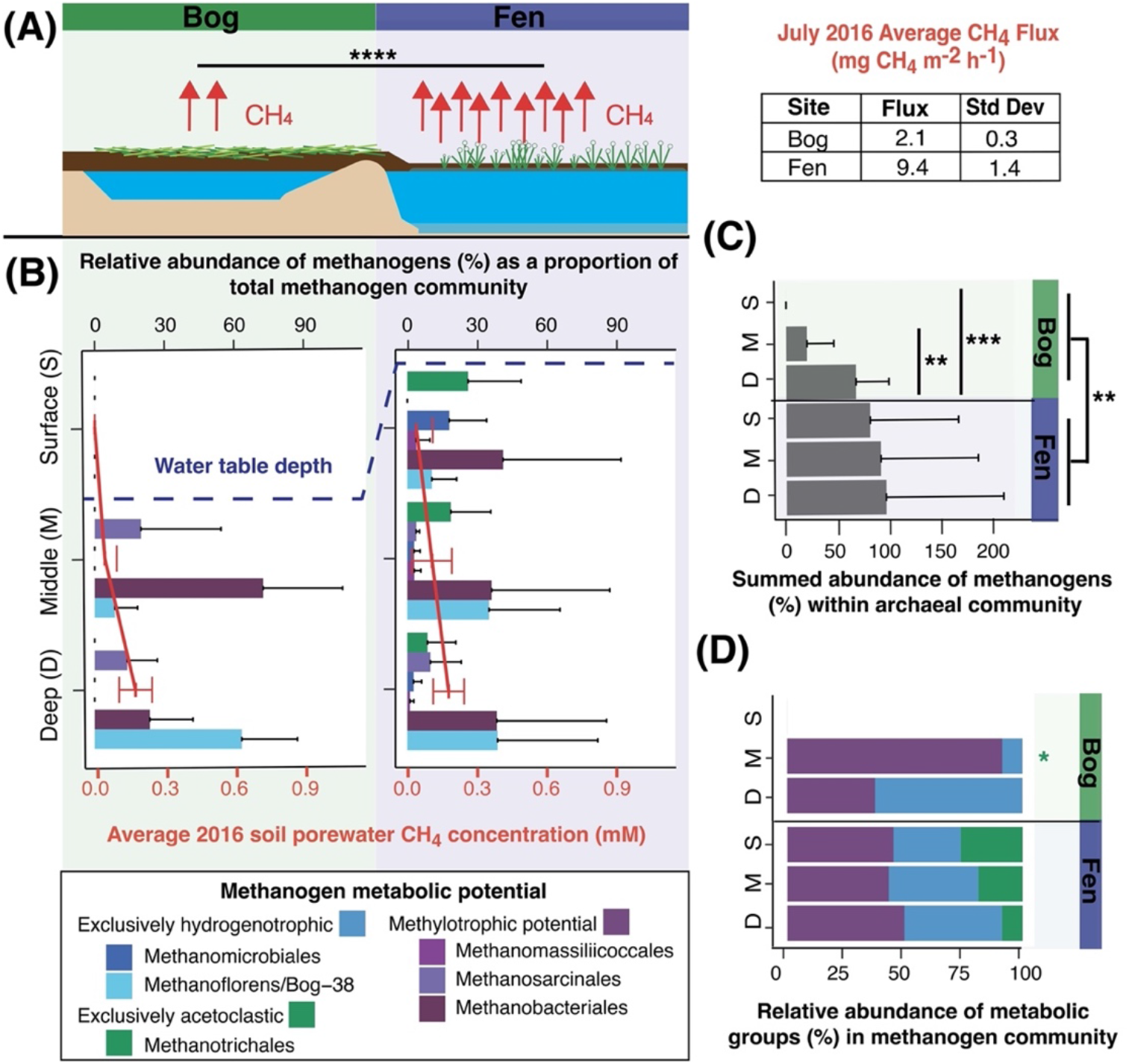
Methanogen abundance and methane flux increase significantly across the Stordalen Mire permafrost thaw gradient in July of 2016. (A) Cartoon showing the structure of the methanogenic habitats in Stordalen Mire. Red arrows represent average methane flux from July 2016, with actual values shown in the table on the right. Flux from the fen is significantly higher than from the bog as per Tukey’s HSD (p-adjusted <0.0001). (B) Bar chart showing the site and depth stratified metagenome-based relative abundance of methanogenic orders within the methanogen community, colored by inferred metabolic potential. The dashed line represents the average water table depth at the time of field sampling. The overlayed red line plot shows the soil porewater methane concentrations from July 2016. Error bars represent one standard deviation for both plot types. The palsa habitat (not shown) showed near negligible production, with values for methanogen relative abundance below detection. (C) Summed relative abundance of all methanogens within the archaeal fraction of the soil microbiota. Error bars represent one standard deviation (and x-axis extends beyond 100% due to error bars). Significant differences were seen within the bog via Tukey’s HSD between the middle and deep (p-adjusted<0.01) and the surface and deep (p-adjusted<0.001). Further, a significant difference in the overall site abundance of methanogens between the fen and bog was found (p-adjusted<0.01). (D) Summed relative abundance of metabolic groups (methylotrophic orders, hydrogenotrophic orders, acetoclastic orders) within the methanogen community. The relative abundance of acetoclastic methanogens was significantly lower than the methylotrophs in the bog at middle depth (Tukey’s HSD, p-adjusted<0.05); otherwise, no significant differences in abundance of acetoclasts or hydrogenotrophs relative to methylotrophs were noted.

The relative abundance of methanogenic taxa presented here is in agreement with past studies in regard to overall community composition (42). However, prior research from Stordalen Mire emphasized the primary importance of hydrogenotrophs – largely *Methanoflorens* – in the bog, and the appearance of acetoclastic *Methanotrichales* in the fen (41, 42). Our expanded sampling and extended genome-resolved physiological curation of these methanogen MAGs revealed a more metabolically diverse community in both the bog and the fen. Both obligate (*Methanomassiliicoccales*) and assigned facultative (*Methanosarcinales* and *Methanobacteriales*) methylotrophic methanogens were found in both bog and fen habitats. Furthermore, the summed abundance of the hydrogenotrophic orders did not differ significantly from the methylotrophic orders in either habitat or depth (Figure 2D). When considering presence of individual MAGs within these orders, on average 64% of the bog (excluding the drained, unsaturated surface) and 30% of the fen methanogen communities encoded the potential for methylotrophic methanogenesis. Our genomic analyses support representation for methylotrophic methanogenesis in this climatically critical ecosystem, warranting further investigation into the chemistry and expressed physiology supporting this metabolism.

### Stordalen Mire peat contains methylotrophic substrates, especially methanol

To further characterize the likelihood of methylotrophy, we next analyzed peat water extracts to detect possible methanogenic substrates. Quantitative nuclear magnetic resonance (NMR) analysis identified 29 such metabolites (Table S4), including classical methanogenic substrates like acetate and formate, as well as the methylated oxygen compound methanol (Figure 3A). The methylated amines glycine betaine and choline were also detected, but only in the non-methanogenic palsa (Figure 3A). Liquid chromatography tandem mass spectrometry (LC MS/MS) also identified methylated compounds as present across habitats (Table S4, Figures S4 & S5) including 4 methylated amines and 4 methylated oxygen compounds (Figure 3B). Of these compounds, only 3 (glycine betaine, choline, and syringate) are recognized as known substrates for this metabolism. Most of the remainder are small derivatives of known substrates (e.g., acylated methylated amines or their stereoisomers) or chemical species with structural homology to known substrates (e.g., apocynin). Notably, trigonelline is chemically distinct and could represent an as-yet unknown aromatic methylamine to support this metabolism.

**Figure 3.**
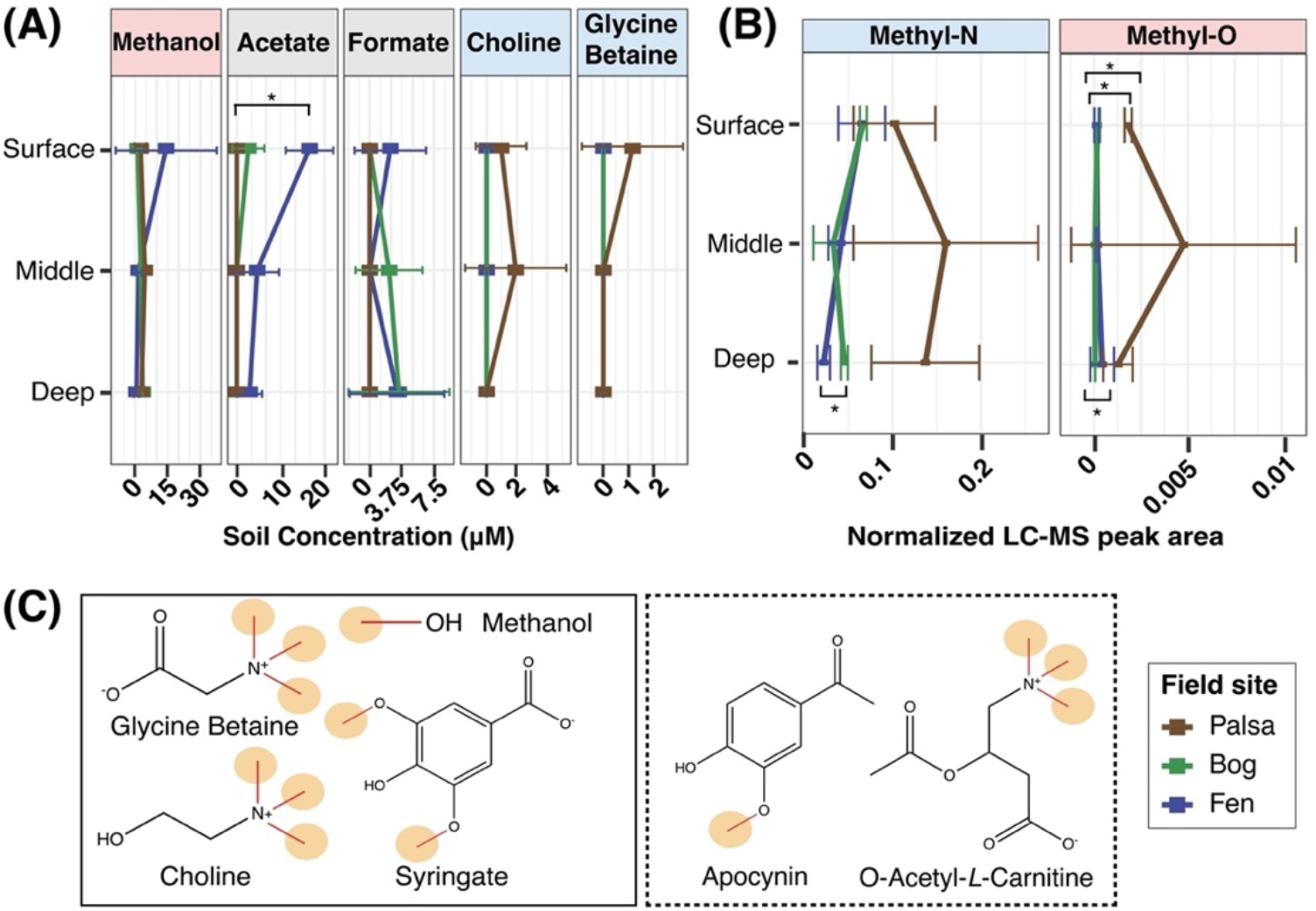
Diverse methylated metabolites are present across Stordalen Mire. (A) Known methanogenic precursors found in peat water extracts as detected by NMR. Points represent average concentrations, with error bars representing one standard deviation. Significant differences were assessed using Tukey’s HSD, with only acetate found to be significantly enriched in the fen compared to the palsa (p-adjusted<0.05). (B) The summed LC MS/MS peak area for 5 methylated amines and 4 methylated oxygen compounds across habitat and depth, which are implicated as potential methylotrophic substrates (Table S4, Figures S4, S5). Methylated amines were found to be significantly higher in the fen than the bog (Tukey’s HSD, p-adjusted<0.05), and significant difference was observed for methylated oxygen compounds between each pair of sites via Tukey’s HSD (Fen:Bog, p-adjusted<0.05; Palsa:Fen, p-adjusted<0.05; Palsa:Bog, p-adjusted<0.05) (C) Chemical structures of select known (solid box) and here proposed possible (dashed box) substrates identified in Stordalen Mire, with microbially available methyl groups circled in orange.

A relevant consideration when trying to link methylotrophic substrates to ecosystem outputs like CH_4_ is that the number of potentially microbially available methyl groups on different compounds (Figure 3C) may limit stoichiometry of CH_4_ formation (18, 20, 21, 38). For example, one molar equivalent of a tri-methoxylated compound may support the production of three times as much CH_4_ as one molar equivalent of methanol. Yet, the broad availability of methanol found across the Mire (e.g., its detection in each palsa, bog, and fen), which may be ultimately plant derived and therefore continually produced in the site (48), lead us to postulate that methanol might be a primary substrate for methylotrophic methanogenesis *in situ*. In support of this, 83% of all methanogen MAGs here classified as methylotrophic were identified as encoding the gene for the methanol-specific methyltransferase MtaB (Figure 1B).

### Methanogens express methylotrophic genes across the Mire

Following investigation of the genomic and chemical potential for methylotrophic methanogenesis, we queried the expression of the putative methylotrophs using genome-resolved metatranscriptomics (Tables S1 & S2). Here we show all three potentially methylotrophic orders are active *in situ* across habitats and depths (Figure 4A). Gene-resolved expression analysis confirmed that almost all (70%) of the active MAGs in these orders were expressing methylotrophic genes (Figure S6). Notably, these actively methylotrophic MAGs represent on average 100% of the summed activity of the *Methanomassiliicoccales*, 85% of the activity of the *Methanosarcinales*, and 91% of the activity of the *Methanobacteriales* across the bog and fen (Figure S6).

**Figure 4.**
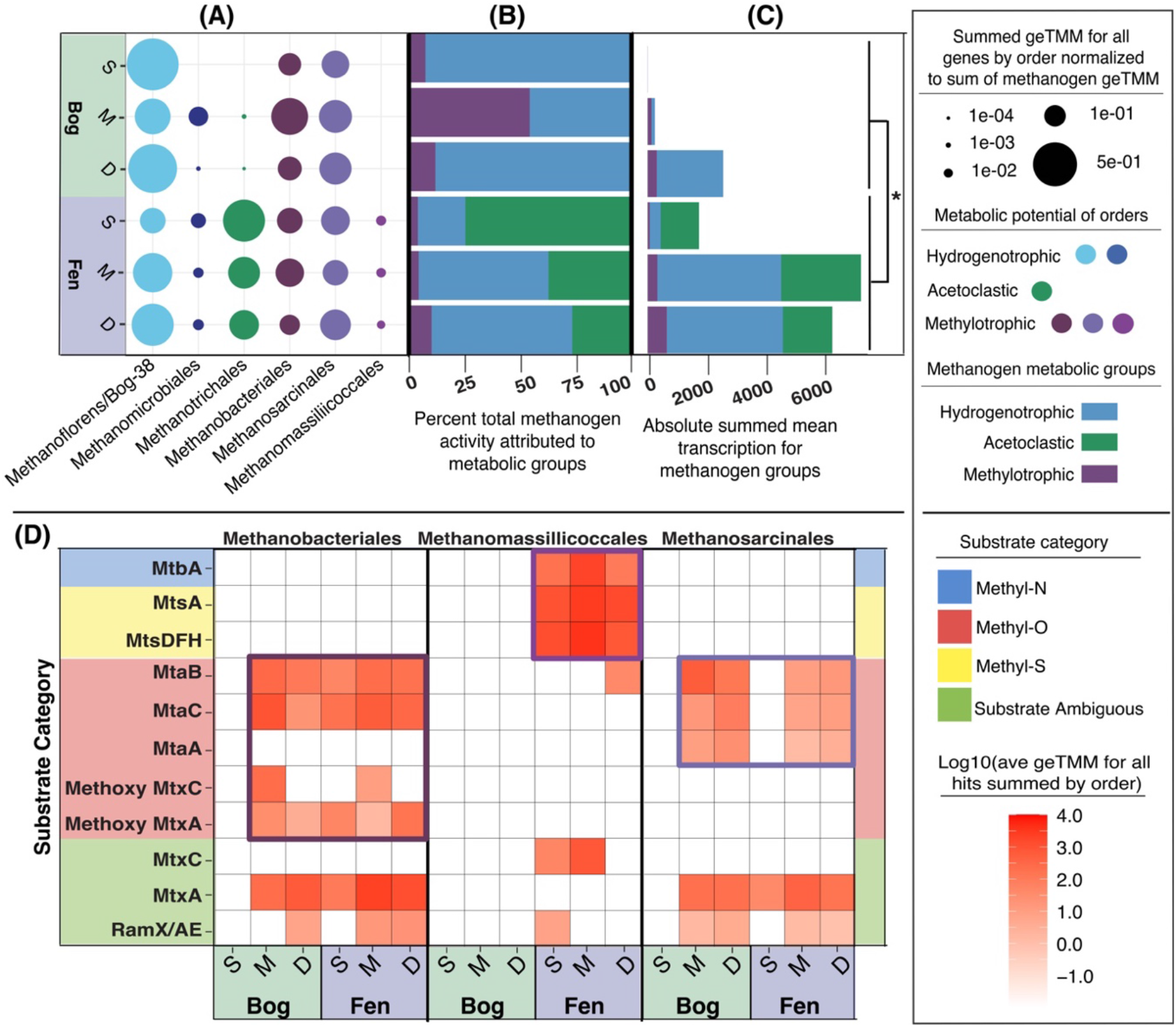
Methanogens with methylotrophic potential are active and expressing genes for methylotrophy across the Mire. (A) Summed relative transcriptional activity of methanogen orders across the Mire within the methanogen community, calculated as averaged geTMM values for all methanogen-expressed genes summed at the order level and normalized to the total sum of all methanogen-expressed genes. (B) Bar chart showing the percent of the total methanogen activity at each depth within the bog and the fen attributable to metabolic groups of hydrogenotrophs, acetoclasts, and methylotrophs. (C) Bar chart showing the absolute summed mean transcription of methanogen metabolic groups across the bog and fen. Total transcription is significantly higher in the fen than bog (Tukey’s HSD, p-adjusted <0.05) but no significant intra-habitat differences were seen between activity of individual metabolic groups. (D) Specific expression of methylotrophic genes by three methanogenic orders across the Mire. geTMM values for expressed methylotrophic methyltransferase genes averaged and normalized to MAG relative abundance within metatranscriptomes and plotted across depth profiles within the bog and fen. Expressed genes are categorized by inferred substrate category. Purple boxes are used to highlight the apparent primary substrate-specific genes expressed by each order. Evidence for active methylotrophic methanogenesis is presented across the bog and fen, at every depth except the bog surface (which is likely the most oxygenated field compartment of the six represented here).

In total, methylotrophic orders accounted for between 5-10% of the total methanogen transcription in the fen, and a broader 7-54% in the bog (Figure 4B) with the greatest proportion of active methylotrophic orders observed in the bog at middle depth. Consistent with our CH_4_ flux data, overall methanogen activity increased significantly between the bog and the fen (Figure 4C). Unlike the acetoclastic methanogens which had significantly increased activity in the fen, no such significant habitat pattern was seen for the methylotrophs. However, per the increased total methanogen activity in the fen relative to the bog, the absolute activity of methylotrophs in the fen, though a smaller average proportion of the methanogen community than in the bog, may not represent an actual decrease in activity with thaw (Figure 4C). Our findings indicate that methylotrophic methanogens play an active role within the Stordalen methanogen community, and thus contribute to the CH_4_ cycle *in situ*.

Our gene expression data was used to refine the substrate usage patterns for these methylotrophic lineages (Figure 4D). For the *Methanomassiliicoccales*, our metatranscript data suggest that methylated sulfides, and possibly methylated amines, are more likely substrates than methylated oxygen substrates, due the limited expression of methanol or methoxy genes. On the other hand, the facultative methylotrophs, *Methanosarcinales* and *Methanobacteriales* exclusively had potential for methylated oxygen usage. Gene expression data supported the use of methylated oxygen compounds, especially methanol, across the bog and fen by these two lineages. It should be noted however that members of all three orders were found to express substrate ambiguous methylotrophic genes, which are not here used to assign functional substrate profiles.

Taken together, our combined metagenomic, metatranscriptomic, and metabolomic data demonstrate active methylotrophic methanogenesis across Stordalen Mire using field-relevant substrates by a sizeable fraction of the native methanogen community. Our expression data also hints at methylotrophic niche partitioning that may occur in the fen, with the *Methanomassiliicoccales* showing preference for methylated sulfide substrates, and the *Methanosarcinales* and *Methanobacteriales* preferentially utilizing methylated oxygen substrates. This simultaneously improves on our understanding of the CH_4_ cycle in this climate critical wetland and demonstrates the need for genome-resolved approaches to studying methanogen physiology.

### Anaerobic methylotrophy is encoded and expressed by numerous bacteria in Stordalen Mire

Some anaerobic bacteria employ homologs to the methanogenic three-component methyltransferase systems, where these same methylotrophic substrates support growth as sources of carbon and/or energy (30, 31, 33–38). Investigation of the methylotrophic potential among the bacterial component of the soil microbiota in Stordalen revealed >1700 MAGs from 19 bacterial phyla encoded thousands of *mtxB* genes (Figure 5A, Table S6). These genes were predominantly inferred to be specific for methylated amines, methanol, and methoxylated compounds. However, some MAGs encoded homologs of methylated sulfide-dependent methyltransferases (Figure 5A), representing to our knowledge the first environmental identification of bacterial methylated sulfide methyltransferase systems. Overall, these data are in good agreement with, and expand upon, the findings of Ticak et al. (30) and Creighbaum et al. (20) on the broad phylogenetic diversity of methylamine-dependent *mtxB* genes extending past solely methanogenic archaea and acetogenic bacteria.

**Figure 5.**
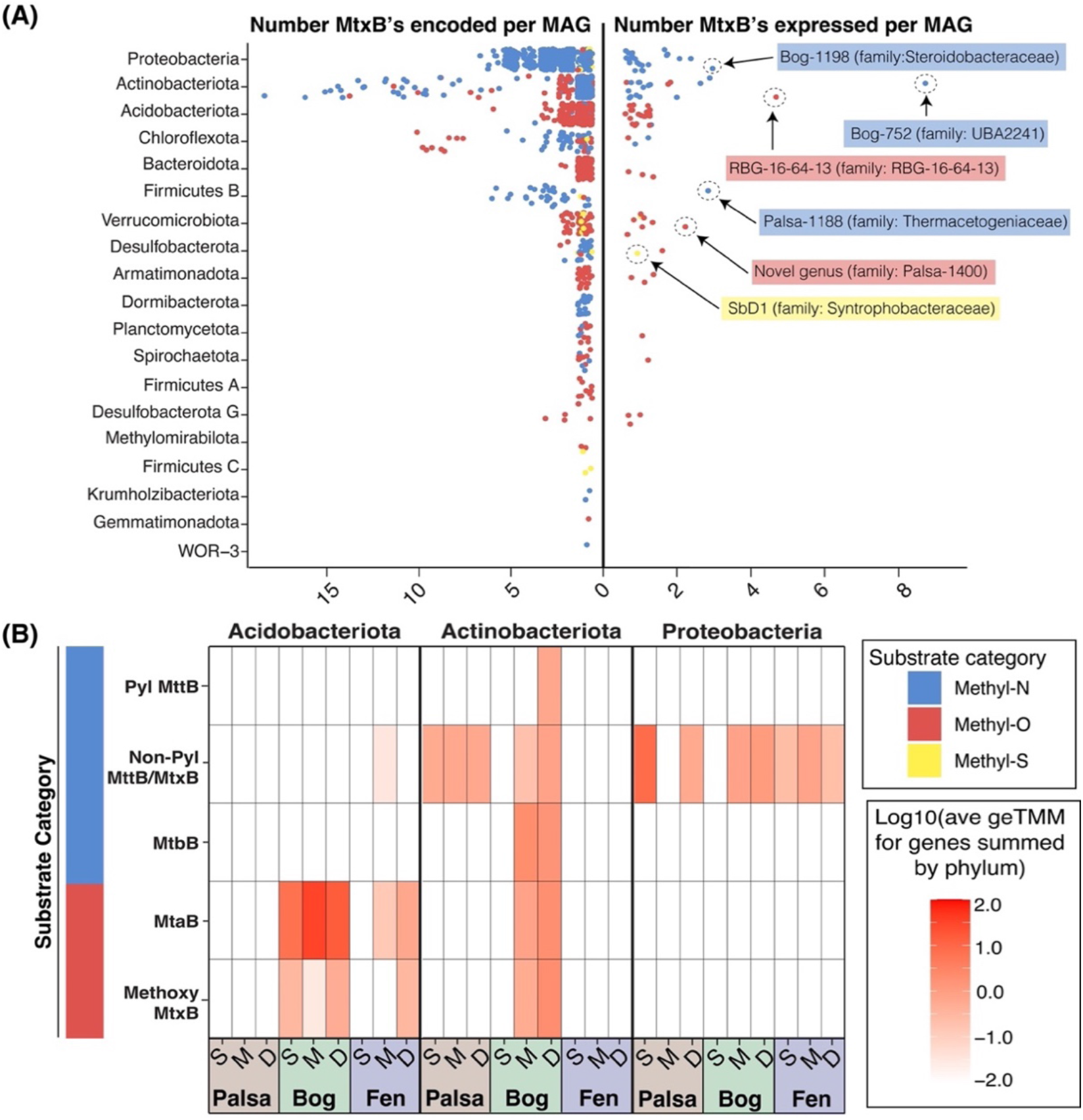
Anaerobic methylotrophic metabolism extends to the bacterial component of the soil microbiota in Stordalen Mire. (A) Plots showing the number of *mtxB* genes encoded (left) and expressed (right) per MAG within bacterial phyla, colored by inferred substrate specificity, in Stordalen Mire. Approximately 1700 bacterial MAGs spanning 19 phyla encode *mtxB* genes, of which 88 from 12 distinct phyla were found to be actively expressing these genes. The genus (and family) of some active methylotrophic bacteria is shown on the right plot to demonstrate the observed taxonomic diversity of the metabolism in Stordalen. (B) Specific expression of identified *mtxB* genes by the three phyla found to include the greatest number of putatively methylotrophic bacteria in the Mire. Bacterial methylotrophic gene expression is evident across the entirety of Stordalen Mire, in both the methanogenic and non-methanogenic habitats.

Of the 19 bacterial phyla here found to encode *mtxB* genes, members of 12 phyla were found to express these genes (Figure 5A), suggesting active bacterial methylotrophy within the Mire. The *Proteobacteria*, *Acidobacteriota*, and *Actinobacteriota* appeared to be the primary methylotrophic bacterial phyla in the site (Figure 5A-5B), with members of genera *Bog-752*, *RBG-16-64-13*, and *Bog-1198* being their most actively methylotrophic representatives. The *Acidobacteriota* exclusively expressed genes for the demethylation of methanol (*mtaB*) and methoxylated compounds across the bog and fen. Members of the *Proteobacteria* expressed *mttB* homologs lacking the unique pyrrolysine residue (non-Pyl) which are known to be specific for quaternary methylated amines such as choline and glycine betaine (30). Notably, members of this phyla expressed these genes in the palsa, where NMR revealed the presence of these compounds (Figure 3A) and our multi-omics failed to detect presence or activity of methanogens. Lastly, the *Actinobacteria* were the most versatile, with expression of genes for demethylation of both methylated amines and methylated oxygen compounds. Interestingly, some MAGs within the *Actinobacteriota* were found to express pyrrolysine-encoding *mttB* and *mtbB* genes, which are specific for tri- and dimethylamine respectively (49). Our findings illustrate new insights into the bacterial transformation of methylated substrates, both phylogenetically showing activity of diverse lineages, and also indicating new substrates for bacterial methylotrophs.

## Discussion

The aim of this study was to employ a genome-resolved approach to query potential for methane-relevant anaerobic methylotrophy, catalyzed by corrinoid-dependent methyltransferase systems, among the microbiota in a thawing permafrost peatland. Here, paired metagenomic and metatranscriptomic data in conjunction with metabolite analyses revealed substantial evidence for metabolism of methylated compounds by members of the soil microbiota (Figure 6). We classified nearly 20% of all Stordalen-derived methanogen MAGs as methylotrophic and further support their activity in saturated soils in the bog and fen using integrated metabolite and metatranscriptomic evidence. Moreover, we extend the role of methylotrophy to the bacterial members of the community, providing a framework for how this metabolism could further compete with or support methanogenesis in a more indirect fashion (Figure 6). Beyond Stordalen, our field data may reciprocally broaden knowledge of the metabolism itself with the identification of wetland-relevant potential novel substrates, plus underappreciated microbial crossfeeding and competition, explored below, which warrant further physiological experimentation.

**Figure 6.**
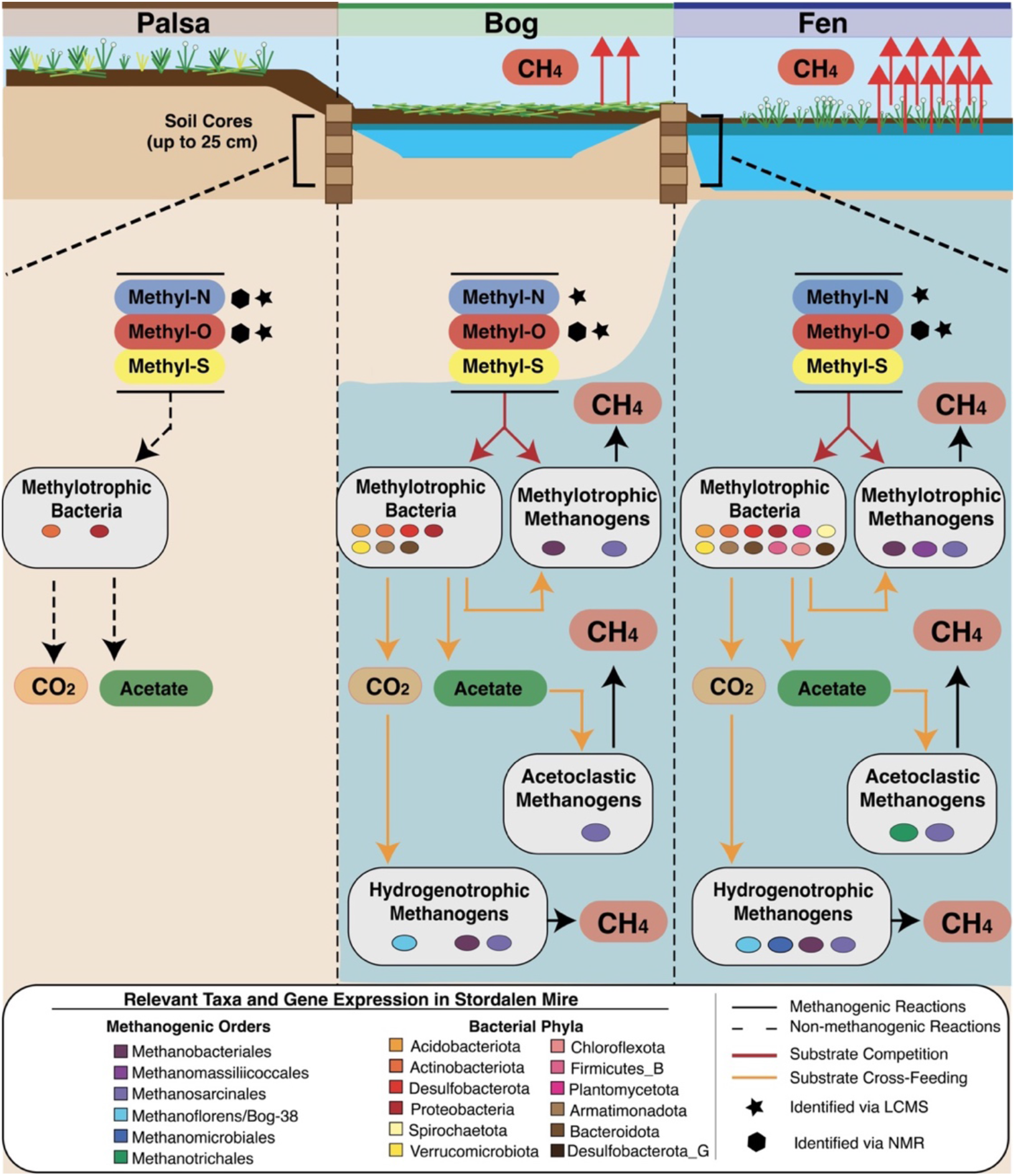
The anaerobic methylotrophic network in Stordalen Mire. Metatranscriptome informed conceptual model summarizing the complexity of the microbial anaerobic methylotrophic food web in Stordalen Mire. Vertical dashed lines separate the palsa, bog, and fen, and the blue background is intended to represent the water table depth within sampled soil cores across habitats relative to microbial metabolic activity. Solid arrows represent metabolic reactions which can lead to the production of CH_4_ either directly or indirectly, while dashed arrows represent reactions not expected to result in CH_4_ production. Red arrows reflect substrate competition, while yellow arrows reflect cross feeding of different metabolic groups. All represented bacterial taxa on the figure include MAGs found to express methylotrophic methyltransferase genes (*mtxB*) in Stordalen Mire. Substrate categories identified within each habitat per metabolite data are represented by stars (LCMS-identified) and hexagons (NMR-identified). Note, the metabolic versatility of the facultatively methylotrophic *Methanosarcinales* and *Methanobacteriales* is represented by their dual inclusion in multiple methanogenic metabolic groups.

### Methylotrophic methanogenesis is encoded by much of the native methanogen community

Though the historical paradigm of environmental methanogenesis has suggested methylotrophy to be a niche metabolism of limited relevance across terrestrial wetlands (50–54), recent studies have countered this idea, increasing both the taxonomic (9–13, 55, 56) and geographic (14–17) footprint of the metabolism. However, a notable challenge remains in profiling environmental methylotrophy. Namely, biochemical discoveries of known relevant genes and substrates are not translated accurately to databases used to functionally annotate genomes or metagenomes (e.g., KEGG). For example, four functionally unique quaternary amine dependent methyltransferases have been biochemically characterized and shown to demethylate distinct substrates (30, 33, 35, 38), but only one of these is present in the KEGG database (30). As a result, the accurate functional assignment of these genes often requires paired non-homology-based methods including gene synteny (19, 30, 33, 38, 57), unique residues (49, 57), and phylogeny (57) to confirm gene functional assignments.

Consequently, holistic surveys of environmental methanogenesis that lack manual methylotrophic curation only screen for a limited number of best-known genes, and thus likely underestimate the overall potential for and diversity of the metabolism. Our annotation strategy to profile methylotrophy in Stordalen Mire did not rely solely on annotation from these databases. For example, via database annotations only, we would have missed most of the *Methanomassiliicoccales*-encoded methylated sulfur use potential, as well as the potential for methanogenesis from methoxylated substrates from any lineage.

Of the methanogenic orders identified here as having potential to catalyze methylotrophy, only the *Methanomassiliicoccales* appear to be functioning as obligate methylotrophs, likely using these substrates dependent on hydrogen as an electron donor (7). Studies have demonstrated that these methanogens are cosmopolitan across water inundated soils and sediments (14, 58–60). In contrast to prior metagenomic studies (14, 58) we did not infer that methanol was the preferential substrate for this lineage. Instead, our metranscriptomic findings indicated preferential utilization of methylated sulfides. While some representative *Methanomassiliicoccales* isolates have been shown to encode the dimethyl sulfide specific *mtsAB* methyltransferase genes (7, 61), no isolates to our knowledge have been shown to grow with methyl sulfides directly (7). Markedly, the Stordalen MAGs were not only found to encode the dimethyl sulfide specific *mtsA*, but also homologs of the tri-functional *mtsD/mtsF/mtsH* genes (23). These genes may confer ability to use a broader suite of methylated sulfide substrates than *mtsAB* alone, which would expand the physiological potential of the order, as well as knowledge of its metabolic niche within wetlands.

Beyond the *Methanomassiliicoccales*, we expanded the observation of methylotrophy within this site to include members of the *Methanosarcinales* and *Methanobacteriales*, identifying these lineages as primarily methanol-utilizing facultative methylotrophs. Both orders have been identified as members of methanogenic communities across global wetlands (60, 62–64). However, the *Methanobacteriales* are typically classified solely as hydrogenotrophs (60, 63), whereas the *Methanosarcinales* are inconsistently labeled as acetoclasts (62), acetoclastic and hydrogenotrophic (60), or metabolically versatile (e.g., capable of all modes of methanogenesis) (14, 63). Prior, two *Methanobacterium* isolates (24, 46) – one of which is from a permafrost system (46) – have been shown to perform hydrogen-dependent methylotrophy using methanol and/or methylamines. Meanwhile, the *Methanosarcinales* are known via biochemical and physiological work to perform all modes of methanogenesis, including methylotrophy from diverse substrates such as methanol (23, 45, 47). Vanwonterghem et al. (9) demonstrated methylotrophic potential for numerous genomes within both orders, including methanol use potential among other wetland (65, 66) and permafrost (67, 68) relevant members. Thus, while both orders are known residents of wetlands, we propose that their classical niches underrepresent their true physiological potential across wetlands, as per their observed gene expression in Stordalen, and potential isolated from other sites.

Considering especially the here identified role for facultative methylotrophs within Stordalen Mire, experimentation is needed to resolve the question of the comparative kinetics of – and thus overall production from - discrete pathways of methanogenesis. This analysis is relevant both at the methanogen community level, but also at the level of a single organism like members of the *Methanosarcina*, where we observed co-expression of acetoclastic and methylotrophic genes in the bog from a single methanogen genome (Table S2). Still, the data presented here imply a more important role for methylotrophy in Stordalen Mire than previously understood. Of 36 methanogen MAGs identified as being active in this dataset, 10 expressed genes for methylotrophy and we confirmed the presence of metabolites to support some of these dominant pathways (Figure 6). Regardless of the kinetics, the widespread footprint of this metabolism across the methanogen community precludes dismissal of its consideration at the ecosystem level.

### Bacterial methylotrophy is a cryptic part of the carbon cycle in wetlands

Though an important role is here supported for methylotrophic methanogens in Stordalen Mire, the metabolism is not limited to the archaeal community. From the literature, it is recognized that anaerobic bacteria, especially certain acetogens, perform methylotrophy via corrinoid-dependent methyltransferase systems feeding into the Wood-Ljungdahl pathway (30, 31, 33–38). Here, we observed expression of *mtxB* genes by bacteria across the entirety of the Mire at nearly all depths, including the drier and undoubtably more oxygenated palsa. While these genes are inferred to be involved exclusively in an obligately anaerobic metabolism, active anaerobes have been identified in oxygenated surface habitats in other locations (69, 70). NMR and metatranscriptomic data support active bacterial methylated amine dependent methylotrophy in the palsa. While not likely supporting methanogens in this habitat, this bacterial metabolism could be a source of CO_2_ (30) an important greenhouse gas.

Overall, these data suggest that bacterial methylotrophy is active across the Mire. In the CH_4_ emitting bog and fen we have considered the ways bacterial methylotrophy could impact the CH_4_ cycle. For instance, while bacterial methylotrophy does not directly produce CH_4_, it may produce acetate (30, 33, 35, 35), and CO_2_ (30), which are substrates for acetoclastic and hydrogenotrophic methanogens respectively. It is also possible that methylotrophic bacteria could cleave quaternary methylated amines (e.g., choline) into smaller methylated amines (trimethylamine) to fuel methylotrophic methanogenesis (71, 72). Methylotrophic bacteria are active in the same habitats and depths as the three metabolic groups of methanogens (Figure S7A), and notably a strong positive correlation was identified between the summed transcriptional activity of methylotrophic methanogens and methylotrophic bacteria in the fen (r^2^ = 0.87) (Figure S7B). Alternatively, bacterial methylotrophs could also compete with methanogens for methylotrophic substrates, complicating this understudied component of the microbial carbon food web. For example, our gene expression data indicates that the *Methanosarcinales* and *Methanobacteriales* (Figure 4D) compete for methanol with the *Acidobacteriota* (Figure 5B). Taken in total, the apparent complexity of the methylotrophic metabolic network in wetlands warrants future experimental work to better resolve the relevance of these metabolisms to wetland methanogenesis, and the terrestrial global carbon cycle. While our data are specific to Stordalen Mire, we can envision extending this model across wetlands per the growing recognition of the importance of methylotrophy across habitats (14–16).

### Conclusions and future needs

Our study advances the growing recognition of the complexity and ecological relevance of methylotrophy, and highlights the power of large-scale field datasets to illuminate its biochemical diversity, phylogenetic extent, and ecological drivers. The methylotrophic metabolic network is increasingly implicated – here and in other climate-relevant ecosystems (14–17) – in impacting atmospheric CH_4_ emissions, especially in a warming climate (17, 73). This expanded knowledge of a widespread metabolism contributing to CH_4_ dynamics in wetlands is essential to improve model-based predictions of wetland contributions to the global CH_4_ budget (75). Process-scale biogeochemical models (like *ecosys* (8)) do not currently account for methylotrophic methanogenesis, representing only acetoclastic and hydrogenotrophic pathways (8, 76) – a likely consequential misrepresentation since methylotrophy dramatically increases the potential route of fixed carbon to CH_4_ production, and its enzymes have distinct kinetics and constraints. This work also provides the field-relevant targets for *in vitro* studies of methyltransferase systems, which are needed to determine field relevant key kinetic parameters, especially since the so far kinetically characterized methylotrophic methyltransferases do not well represent known wetland methanogen communities. Quantifying the methylotrophic contribution of methanogenesis in wetlands – the largest natural biogenic CH_4_ source (67) – likely via isotopically labeled substrate experiments (77), is an essential next step in constraining its addition to predictive models. Together, these integrated approaches will increase biological realism in models and predictions for these and other major CH_4_ -emitting, climate sensitive habitats.

## Methods

### Field site and sample collection

Stordalen Mire (68 22’N, 19 03’E) is a rapidly thawing Artic permafrost peatland near Abisko, Sweden. The mosaic of permafrost land cover includes three primary biologically and chemically distinct habitats that constitute a discontinuous permafrost thaw gradient. The raised, well-drained palsa overlays intact permafrost with an active layer depth of 50-60 cm and is dominated by woody and ericaceous shrubs. The bog site, dominated by *Sphagnum* mosses, is underlain by partially thawed permafrost and a seasonally fluctuating perched water table. Last, the sedge-dominated fen represents fully thawed and saturated permafrost. In July 2016, cores were taken for meta-omic and geochemical analyses, in triplicate using an 11-cm diameter push corer from each palsa, bog, and fen in areas adjacent to the *in situ* gas flux measurement autochamber system. Cores were sub-sectioned in the field at three depths: surface (1-4cm), middle (10-14cm), and deep (20-24cm). Subsections were split based on end-use; for nucleic acid extraction, 4 ml of field-saturated peat were added to 2.5 volumes of Lifeguard buffer (Qiagen, Maryland, USA), transferred out of the field on ice in a cooler, and frozen at -80°C until extraction. A second split – a portion of which was used for metabolomics – was placed in a 50 ml Falcon tube without buffer, flash frozen in liquid nitrogen, transferred out of the field on ice in a cooler, and stored at -80°C until processing.

### Methane field measurements

To determine soil porewater CH_4_ concentrations, prior to coring at each site, porewater was collected with a perforated stainless-steel piezometer inserted into the peat and extracted with an airtight syringe. No porewater was obtained from sites (palsa) or depths (in the bog or fen) that were above the water table. Porewater samples were filtered and acidified and stored in evacuated vials until they were brought to atmospheric pressure with helium, and CH_4_ concentration in the equilibrated headspace was measured using a Flame-Ionization-Detector Gas Chromatograph. An extraction efficiency of 0.95 was used to calculate dissolved CH_4_ concentration.

CH_4_ fluxes were measured using a system of 8 automated gas-sampling chambers made of transparent Lexan (n=3 each in the palsa, bog, and fen habitats, with n=2 in the fen prior to 2011). Chambers were initially installed in 2001 (78) and the chamber lids were replaced in 2011 with the larger current design, similar to that described by Bubier et al. 2003 (79). Each chamber covers an area of 0.2 m^2^ (45 cm × 45 cm), with a height ranging from 15-75 cm depending on habitat vegetation. At the palsa and bog sites the chamber base is flush with the ground and the chamber lid (15 cm in height) lifts clear off the base between closures. At the fen site the chamber base is raised 50–60 cm on lexan skirts to accommodate large-stature vegetation. The chambers are instrumented with thermocouples measuring air and surface ground temperature, and water table depth and thaw depth are measured manually 3–5 times per week. The chambers are connected to the gas analysis system, located in an adjacent temperature-controlled cabin, by 3/8” Dekoron tubing through which air is circulated at approximately 2.5 L min^-1^. Each chamber lid is closed once every 3 hours for a period of 8 minutes, with a 5-minute flush period before and after lid closure. Results for multiple years are reported in Holmes et al., 2022 (40).

July 2016 CH_4_ flux data and porewater data are in Table S4; data for field sites including porewater CH_4_ and water table depth can be found at 10.5281/zenodo.7720573, and the July 2016 flux data used in Figure 1 can be found at 10.5281/zenodo.7897922.

### Metagenome Assembly and Binning

To maximize the site-specific MAGs available for this study, they were developed from 3 sources: (i) all those generated from Woodcroft and Singleton et al. (43), (ii) assembly and binning of metagenomes from 2010-2017 field samples, and (iii) all fractions of a stable isotope probing experiment performed on field peat added to labeled plant matter from locally co-occurring species. Stable isotope probing metagenomes were used here purely to add site MAGs, not for analyses of that experiment.

Metagenome read sets from 2010-2017 field samples were trimmed using trimmomatic (v0.36) (80) in the paired-end mode against the TruSeq3-PE illumina adapters with a sliding window of 4 to 15. Trimmed reads were then assembled with SPAdes (v3.12, --meta option enabled) (81) with the default kmer set. Sample 20120700_E3M was too complex to assemble the first k-mer set within our compute limits, so the reads were randomly subsampled to 50% using bbtools (v38.51) (82) reformat.sh and assembled with metaSPAdes (83). The contigs from this sample were then dereplicated before subsequent steps with cdhit (v4.8.1) (84) with the following parameter sets: -c .99 -aS .80 -n 11 -d 0 -g 1. Read-mapping was done from the quality-controlled reads against all samples. Once assembly and read mapping were complete, the bam files and contigs were used as input for binning. An initial bin set was created using UniteM (v0.0.18) (85) with the following options: mb_sensitive, mb_verysensitive, mb_specific, mb_veryspecific, mb2, max40, max107, bs, gm2. Following, UniteM, DAS Tool (v1.1.1) (86), and MetaWRAP (v1.0.6) (87) were used to create ensemble bin sets. Due to the limitation of the MetaWRAP bin_refinement module only accepting three candidate bin sets, MetaBAT2 (88), GroopM2 (89), and MaxBin2 (90) from the initial bin sets were used as input into MetaWRAP. The output of these ensemble tools was then used as input into the same tools (DAS Tool, MetaWRAP, and UniteM) for a second iteration of ensemble binning.

Completion and contamination statistics of the second iteration of ensemble bins were determined using CheckM (v1.0.12) (91) lineage workflow. Bins with at least 70% completion and less than 10% contamination were leveraged to determine a quality score of the three ensemble bin sets for each sample individually. The quality score was calculated as follows: score = completeness - (5 x contamination). The ensemble binning tool with the highest quality score was used as the bin set for that sample. For any samples where the number of bins generated from the first step of UniteM were too large for our computational limits, MetaBAT2 was used alone. Once a candidate bin set was chosen, RefineM (v0.0.24) (92) “outliers” was run using the following parameters: --td_perc 95 --gc_perc 95. All MAGs with at least 70% completion and less than 10% contamination were then manually examined and refined through anvi’o margaret (v5.2) (93) leveraging differential coverage and GC content with hierarchical clustering guiding refinements.

Read sets from the SIP experiment were trimmed identically, though were assembled with both SPAdes (--meta option enabled) (v3.13) with the default kmer set, and with MEGAHIT (v1.1.3) (94) with the default kmer set. Additionally, BFC error (95) correction was performed and assembled with SPAdes (v3.13) with the default kmer set. For all samples that had 2 sets of reads, the largest read set was assembled. Abundance information for each contig was generated using bowtie2 (v2.4.1) (96) by mapping reads from all samples (without T0), within the same habitat, to all contigs, for each assembly. T0 reads were only used on assemblies generated from T0 samples. Each assembly was binned separately into MAGs using MetaBAT2 (v2.12.1).

### Additional Sources

Genomes and assemblies for the stable isotope probing samples were downloaded from JGI on December 7, 2020, and December 4, 2020, respectively. These datasets were generated through the DOE-JGI metagenome annotation pipeline (97).

From these total efforts, a database of 13,290 medium and high-quality (98) MAGs was generated using data from 882 Stordalen Mire field and microcosm metagenomes spanning sampling from 2011-2017. MAGs were annotated using DRAM (v1.3.2) (99). Taxonomy was assigned using GTDB-Tk (v2.1.1 r207) (100). These 13,290 MAGs were dereplicated at 97% identity using dRep (101) into 1,864 representative MAGs with galah (v0.3.0) (102) using the following parameter set: --precluster-ani 90 --ani 97 --precluster-method finch. Accession information for metagenomic reads is provided in Table S1, and the full database of 13,290 MAGs can be downloaded from https://doi.org/10.5281/zenodo.7596016.

### RNA Extraction

DNA and RNA were extracted using the Mobio PowerMax Soil DNA/RNA isolation kit (cat# 12966-10). Sample vials were removed from the -80°C freezer and thawed on ice. Following this, 5-10g of peat materials (preserved in Lifeguard soil preservation solution, Qiagen) was added into bead tubes, and nucleic acids were extracted per manufacturer guidelines without initial addition of beta-mercaptoethanol. Reagents were increased proportionally to maintain the concentration of solutions. An additional ethanol wash of the nucleic acids-bound column was performed to further remove impurities. Resulting washed nucleic acids were eluted with 5 mL RNase-free DI water, and further concentrated via ethanol precipitation overnight, followed by elution in 100 µL of TE buffer. The eluted product was further processed for separation and purification of each DNA and RNA; samples were aliquoted into two 2 mL tubes at a ratio of 1:2. RNase and DNase treatment (Roche) were both performed following manufacturer guidelines, followed by phenol:chloroform purification. Nucleic acids were then ethanol precipitated, and pellets were eluted in TE buffer. Purified DNA and RNA were quantified via Quibit 3.0. All samples were stored at -80°C pending sequencing.

### Metatranscriptome Analysis

Metatranscriptome libraries were prepared for 27 field samples at Ohio State University. Using 10ng RNA for each, rRNA was first depleted using the QIAseq FastSelect -5S/16S/23S (Qiagen) kit per the protocol with some modifications: probes for both plants and yeast were added, and only one-third of the probe volumes were used. Next, the TruSeq Stranded Library Preparation kit (Illumina) was used to prepare the sequencing library. Libraries were sequenced on an Illumina NovaSeq 6000 system at the Genomics Core at the University of Colorado Anschutz Medical Campus.

Raw metatranscriptome reads were quality trimmed and adapters were removed using bbduk (82) with the following flags: k=23 mink=11 hdist=1 qtrim=rl trmiq=20 minlength=75. Reads were filtered with rqcfilter2 using the following flags: jni=t rna=t trimfragadapter=t qtrim=r trimq=0 maxns=1 maq=10 minlen=51 mlf=0.33 phix=t removeribo=t removehuman=t removedog=t removecat=t removemouse=t khist=t removemicrobes=t mtst=t sketch kapa=t clumpify=t tmpdir=null barcodefilter=f trimpolyg=5. Trimmed filtered reads were mapped using bowtie2 (96) against the dereplicated MAG database (n=1864 MAGs) with the following settings: -D 10 -R 2 -N 1 -L 22 -I S,0,2.50. The resulting SAM file was converted to a BAM with samtools (103) and then filtered using the reformat.sh script from bbtools (82) with the following settings: idfilter=0.95 pairedonly=t primaryonly=t. Mapped reads were counted with htseq-count (104) with the flags: -a 0 -t CDS -I ID –stranded=reverse. Last, read counts were filtered to remove counts <5, and were converted to geTMM values (105) in R. Metatranscriptomic reads are available on NCBI, with accession numbers reported in Table S1.

### Initial profiling of methanogen physiology across Stordalen Mire MAGs

To first query the taxonomically presumed methanogen MAGs for methanogenic potential (Figure S2A), the DRAM output for 367 MAGs spanning six orders was analyzed for the presence of a variety of methanogenesis genes, including those encoding the Mcr complex and the heterodisulfide reductase complex (Hdr). KOs for all genes screened here are listed in Table S2. Of 367 MAGs screened, 82% encoded part of the Mcr complex but 99% encoded either Mcr or Hdr. Only 4 MAGs (2 *Methanoflorens* and 2 *Methanomicrobiales*) lacked genes encoding subunits of either of these protein complexes. However, as these 4 MAGs each encoded most of the Wood-Ljungdahl pathway, subunits of the Mtr complex, and characteristic hydrogenases of hydrogenotrophic methanogens (Figure S2A) – plus belonged to well resolved methanogenic genera/species represented by other Mcr/Hdr-encoding MAGs in this dataset - they were still considered likely methanogens in this study.

Methanogenic MAGs were then screened for indicators (5, 6, 106–109) of specific methanogen physiology (hydrogenotrophic, acetoclastic, methylotrophic) (Figure S2B-C). All 220 *Methanoflorens* MAGs were observed to encode at least part of the Wood-Ljungdahl pathways, most the Mtr complex, plus subunits of hydrogenase complexes (Mvh, Frh) that support their classification as hydrogenotrophic. Similarly, the designated-hydrogenotrophic MAGs within the *Methanomicrobiales* all encoded part of the Wood-Ljungdahl pathway, most of the Mtr complex, and at least one relevant hydrogenase (Eha, Frh, Ech) per MAG.

The MAGs classified as members of the *Methanotrichales*, obligately acetoclastic per literature (109), were found to encode genes relevant for acetoclastic methanogenesis including the acetyl-coA synthase/carbon monoxide dehydrogenase complex (ACS-CODH) and acetyl-coA synthetase. Meanwhile the *Methanosarcinales* – often considered acetoclastic in wetland literature, but biochemically known to catalyze all modes of methanogenesis – were all found to encode acetate kinase and phosphate acetyltransferase plus the ACS-CODH (suggesting acetoclastic potential), most the Mtr complex, as well as the Wood-Ljungdahl pathway and hydrogenotrophic hydrogenases (including Vht, Ech, Frh). Multiple MAGs were found via DRAM to encode methylotrophic genes, however a detailed discussion of a more in-depth curation of methylotrophy follows below.

The *Methanobacteriales* were all found to encode the Wood-Ljungdahl pathway as well as hydrogenotrophy-relevant hydrogenases (Eha, Ehb, Frh, Mvh), though many were found to lack the Mtr complex. Regardless, these MAGs were considered capable of hydrogenotrophy. Additionally, many were noted to encode methylotrophic genes. Last, the *Methanomassillicoccales* MAGs were found to lack the Wood-Ljungdahl pathway but to encode at least some relevant hydrogenases (Mch, Ech) for hydrogen-dependent methylotrophy, leading to their designation as obligate methylotrophs.

Following this DRAM-based analysis, the methylotrophic potential of these MAGs was further manually curated to better assess the substrate-specific potential for methylotrophy across these MAGs.

### Further profiling of methylotrophy across Stordalen Mire MAGs

To better resolve the potential of the Stordalen methanogens for methylotrophy, the MAGs were searched via BLAST-P using a FASTA reference file (Figure S2C, Table S3) of 53 well-characterized methylotrophic gene types (20 *mtxB* genes, 16 *mtxC* genes, 10 *mtxA* genes, and 7 *ram* genes). The BLAST-P output (Table S2) was limited to include only hits with a bitscore >60 from MAGs found to encode homologs of *mtxB* genes. Genes passing this threshold were phylogenetically analyzed using ProtPipeliner to build RaxML trees (https://github.com/TheWrightonLab/Protpipeliner/blob/master/protpipeliner.py) relative to reference genes including those used in the BLAST-P search, plus other homologous sequences derived from UniProt from physiologically characterized methylotrophic methanogens and acetogens (Table S3). Newick trees are available https://doi.org/10.5281/zenodo.7864933. Gene trees were built for *mtxB* (127 genes), *mtxA* (168 genes), *mtxC* (190 genes), *ramX* (100 genes), and methylated sulfur genes (36 genes). Trees were visually inspected in iTOL (110), and tree placement was used to confirm or refine the specific identification of genes (Table S2). In some cases, genes were only ambiguously identified as methylotrophy-relevant but substrate-nonspecific *mtxA* or *mtxC*. RamX proteins are known to be promiscuous activating enzymes across corrinoid proteins (111), and so activase-encoding genes were identified as evidence for methylotrophy in conjunction with *mtxBCA* genes, but not used to infer substrate specificity for MAGs.

Eighty-five of the 86 MAGs belonging to the *Methanobacteriales*, *Methanosarcinales* and the *Methanomassiliicoccales* were found to encode genes for methylotrophy and used throughout the remaining analyses of methylotrophic methanogenesis. However, MAGs were conservatively defined as methylotrophic only if found to encode genes for at least two types of the three core members of a methyltransferase system directly involved in substrate demethylation (MtxB, MtxC, MtxA), one of which had to be the substrate demethylating *mtxB*. In the case of methyl sulfide metabolism, MAGs were screened for having single genes encoding any of the tri functional MtsDFH proteins, or at least one of *mtsAB* (See Figure 1C, Figure S2). To determine if methylotrophy was being expressed within the MAGs, identified methylotrophic genes were mined from paired metatranscriptomic data (Table S2). Analogous rules were used to label MAGs as active methylotrophs only if found to be expressing the majority of an identified methyltransferase system including an *mtxB* gene. To determine overall order-level methanogen field activity, average (n=3) geTMM values for all methanogen-expressed genes were mined from the data (Table S2). Here, the *Methanomassiliicoccales* were identified as obligate likely hydrogen-dependent methylotrophs, while the *Methanobacteriales* and *Methanosarcinales* were classified as facultative methylotrophs.

Next, to query the bacterial community for methylotrophy genes, a similar BLAST-P approach was used to query the 12,868 bacterial MAGs just for *mtxB* genes – considered here the best single marker gene for methylotrophy - with a minimum bitscore >200 (Table S6). A subset of the same (Table S3) FASTA reference file was used, limited to include only the *mtxB* genes. Bacterial MAGs were only screened for the highly substrate-specific *mtxB* as the best marker gene of methylotrophy, and to reduce potential nonspecific hits to other methylotrophy genes (e.g., other bacterial cobalamin-binding proteins). Identified genes were parsed from metatranscriptomes to determine active bacterial methylotrophy. Phylogenetic analysis using ProtPipeliner (Newick trees available at https://doi.org/10.5281/zenodo.7864933) to confirm the substrate-resolved identity of *mtxB* genes was performed only for those encoded and expressed by the *Acidobacteriota, Actinobacteriota*, and the *Proteobacteria* represented in Figure 7B. To distinguish between bacterial *mttB* genes encoding and lacking the unique pyrrolysine (Pyl) residue, sequences were analyzed using Geneious 2023.0.1 (www.geneious.com) to look for gene truncation due to the amber codon encoding Pyl. Those found to be truncated were identified as “Pyl-MttB” (known to be specific to tri/di/monomethylamine (49)), and those found to lack said truncation were identified as “Non-Pyl MttB/MtxB” (known to be specific for quaternary amines (30)) (Figure 5B).

### MAG Metagenome Relative Abundance Determination

To determine metagenome abundances of the 97% dereplicated MAG set, we first mapped trimmed metagenome reads to the MAG set using bowtie2 (v2.4.5) (96) using the following settings: -D 10 -R 2 -N 1 -L 22 -i S,0,2.50. The output SAM file was converted to a sorted BAM, using samtools (v1.9) (112), and filtered using the reformat.sh script in the bbtools (82) package using: idfilter=0.95 pairedonly=t primaryonly=t. MAG abundance was inferred from this using coverM (v0.6.0). coverM (v0.6.0) genome (https://github.com/wwood/CoverM) was then run using the produced BAM files as input, to calculate the coverage of the dereplicated MAGs within the field, to permit calculation of methanogen relative abundance. CoverM was run with the following flags: coverm genome --proper-pairs-only -x fna –min-read-percent-identity-pair 0.95 –min-read-alignment-percent-pair 0.75 -m trimmed-means –trim-min 0.1 – trim-max 0.9. MAG relative abundances were calculated using the output in R. A final table containing the relative abundance of all MAGs within the 97% dereplicated set, plus the abundances of methanogen MAGs normalized to both the total summed abundance of all archaea and all methanogens is in Table S5.

### Phylogenomic analysis of native methanogen community

Phylogenomic analysis of the 367 Stordalen Mire methanogen MAGs (Table S1) was performed using GTDB-Tk v2.1.1 r207 (100) run using the de novo workflow. The alignment was based on 53 concatenated archaeal marker genes, and a GTDB-derived genome from the phylum Undinarchaeota (GCA_002495465.1) was used as an outgroup to root the tree. The generated tree was read and visually modified in R using the ggtree package (113). Newick tree is available at https://doi.org/10.5281/zenodo.7864933.

### Metabolite extraction and LC MS/MS

Water soluble metabolites were extracted from peat by adding 7 mL of autoclaved milliQ water to 1g of wet peat in a sterile 15 mL centrifuge tube. Tubes were vortexed for 30 seconds two times, sonicated for 2 hours at 22°C, and then centrifuged; the resulting 6 mL supernatant served as the water extract. 2 mL of this was aliquoted for LC MS/MS and stored at -80°C until use.

Water-soluble extracted metabolites were thawed at room temperature and centrifuged to remove any resultant particulates. Each sample was divided into 2 1 mL aliquots in 2 mL glass tube vials for hydrophilic interaction liquid chromatography (HILIC) and reverse-phase (RP) liquid chromatography respectively. Both vials were dried down completely on a Vacufuge plus (Eppendorf, USA). Samples for HILIC were resuspended in 50:50 solution of acetonitrile and water. Samples for RP were resuspended in a 20:80 solution of HPLC grade methanol in water.

A Thermo Scientific Vanquish Duo ultra-high performance liquid chromatography system (UHPLC) was used here for liquid chromatography. Extracts were separated with a Waters ACQUITY HSS T3 C18 column for RP separation and a Waters ACQUITY BEH amide column for HILIC separation.

Samples were injected in a 1 μL volume onto the column and eluted as follows: for RP the gradient went from 99% mobile phase A (0.1% formic acid in H_2_O) to 95% mobile phase B (0.1% formic acid in methanol) over 16 minutes. For HILIC the gradient went from 99% mobile phase A (0.1% formic acid, 10 mM ammonium acetate, 90% acetonitrile, 10% H_2_O) to 95% mobile phase B (0.1% formic acid, 10 mM ammonium acetate, 50% acetonitrile, 50% H_2_O). Both columns were run at 45 °C with a flowrate of 300 μL/min.

A Thermo Scientific Orbitrap Exploris 480 was used for spectral data collection with a spray voltage of 3500 V for positive mode (for RP) and 2500 V for negative mode (for HILIC) using the H-ESI source. The ion transfer tube and vaporizer temperature were both 350 °C. Compounds were fragmented using data-dependent MS/MS with HCD collision energies of 20, 40, and 80.

The commercially available Compound Discoverer 3.2 software (Thermo Fisher Scientific) was used to analyze the data using the untargeted metabolomics workflow. Briefly, the spectra were first aligned followed by a peak picking step. Putative elemental compositions of unknown compounds were predicted using the exact mass, isotopic pattern, fine isotopic pattern, and MS/MS data using the built in HighChem Fragmentation Library of reference fragmentation mechanisms. Metabolite annotation was performed first by matching fragmentation scans, retention time and ion mass to an in-house database built using 1200 reference standards. Second, fragmentation scans (MS2) searches in mzCloud were performed, which is a curated database of MSn spectra containing more than 9 million spectra and 20000 compounds.

Third, predicted compositions were obtained based on mass error, matched isotopes, missing number of matched fragments, spectral similarity score (calculated by matching theoretical and measured isotope pattern), matched intensity percentage of the theoretical pattern, the relevant portion of MS, and the MS/MS scan. The mass tolerance used for estimating predicted composition was 5 ppm. Finally, annotation was complemented by searching MS1 scans on different online databases with ChemSpider (using either the exact mass or the predicted formula). Based on the annotation results, metabolites were divided into four categories, and were assigned as either Level 1, Level 2, or Level 3 following the Metabolomics Standards Initiative (114) : (1) full match to in-house database, (2) full match based on mzCloud, predicted composition, and ChemSpider, (3) full match based on predicted composition and Chemspider and (4) annotated by only one method (ChemSpider) with potential annotation being based on mass alone.

The structures of all chemical compounds identified via LC MS/MS (Table S4) in Stordalen Mire were examined only for any rank-1 species including a methylated nitrogen, oxygen, or sulfur atom. Methylated compounds which met this criterion were then compared to known methylotrophic substrates (Figure S3) to look for structural homology. Note, compounds of interest for this study were only identified in the RP, and not the HILIC, data, and thus only the former is of focus in this manuscript. Peak areas for identified compounds were normalized to the sum of all compounds within each individual sample prior to further analysis in R. Boxplots made in R v4.1.1(115) (Figure S4) show the individual averaged (n=3) LC MS/MS peak area for each species over thaw and depth gradients; these averaged peak areas were categorically summed by “methyl-N”/methylated amines” and “methyl-O”/methylated oxygen compounds which is shown in Figure 3B. LC MS/MS data can be found in https://doi.org/10.5281/zenodo.7519815.

### 1H NMR

To identify field-present metabolites in Stordalen, including methanogenic and methylotrophic precursors, we performed ^1^H NMR on aliquots of the same peat extractions prepared for LC MS/MS analyses. Samples (180 µL) were combined with 2,2-dimethyl-2-silapentane-5-sulfonate-d_6_ (DSS-d_6_) in D_2_O (20 µL, 5 mM) and thoroughly mixed prior to transfer to 3 mm NMR tubes. NMR spectra were acquired on a Varian 600 MHz VNMRS spectrometer equipped with a 5 mm triple-resonance (HCN) cold probe at a regulated temperature of 298 K. The 90° ^1^H pulse was calibrated prior to the measurement of each sample. The one-dimensional ^1^H spectra were acquired using a nuclear Overhauser effect spectroscopy (NOESY) pulse sequence with a spectral width of 12 ppm and 512 transients. The NOESY mixing time was 100 ms and the acquisition time was 4 s followed by a relaxation delay of 1.5 s during which presaturation of the water signal was applied. Time domain free induction decays (57472 total points) were zero filled to 131072 total points prior to Fourier transform. Chemical shifts were referenced to the ^1^H methyl signal in DSS-d_6_ at 0 ppm. The 1D ^1^H spectra were manually processed, assigned metabolite identifications, and quantified using Chenomx NMR Suite 8.3. Metabolite identification was based on matching the chemical shift, J-coupling and intensity of experimental signals to compound signals in the Chenomx and custom in-house databases. Quantification was based on fitted metabolite signals relative to the internal standard (DSS-d_6_). Signal to noise ratios (S/N) were measured using MestReNova 14 with the limit of quantification equal to a S/N of 10 and the limit of detection equal to a S/N of 3. Known methanogenic substrates were manually identified from the list of quantitated metabolites, and average (n=3) concentrations were plotted using R v4.1.1(115) shown in Fig 3. Summarized data is available in Table S4, and raw data can be found at https://doi.org/10.5281/zenodo.7519683.

### Data Visualization and Statistics

Data were analyzed and visualized in R v4.1.1 (115, 116) using ggplot2 (117) unless otherwise specified. The superheat package was used to generate the heatmap in Figure 1D. All reported statistical analyses including ANOVA, TukeyHSD, and Pearson Correlation were performed in R using the ggpubr package (118).

### Data Availability

The metagenomes, metatranscriptomes, and metagenome-assembled genomes used in this paper are available at NCBI under BioProjectID PRJNA386568. Table S1 provides individual BioSample numbers. Raw and processed data for all analyses are available in the following Zenodo archives: 97% dereplicated MAG annotations (10.5281/zenodo.7587534), metatranscriptome mapping (10.5281/zenodo.7591900, LC MS/MS (10.5281/zenodo.7519815), NMR (10.5281/zenodo.7519683), July 2016 methane flux data (10.5281/zenodo.7897922), Newick trees for phylogenomic and phylogenetic analyses plus DRAM v1.3.2 annotations for all methanogens and methylotrophic bacteria (10.5281/zenodo.7864933). Note, MAGs have been submitted to NCBI but are still processing through their system; they are being temporarily housed in the Zenodo archive (10.5281/zenodo.7596016); this Zenodo archive will ultimately re-route readers to the final NCBI deposition. Photographs of the Mire used in Figure S1 (file names: AJ_palsa_05.jpg, Eriophorum_eample_06.jpg, Sphagnum_04_(drier).jpg) were all taken by Nicole Raab in July 2016, and retrieved/available from the EMERGE database (emerge-db.asc.ohio-state.edu). Last, the full field metadata sheet for Stordalen Mire published by EMERGE is available at 10.5281/zenodo.7720573.

## Abbreviations

CH_4_: Methane; CO_2_: Carbon Dioxide; MAGs: Metagenome-assembled genome; *mcrA*: Methyl-coenzyme M reductase alpha subunit; NMR: Nuclear magnetic resonance; LC MS/MS: Liquid chromatography-tandem mass spectrometry; MtxB: Substrate:corrinoid methyltransferase; MtxC: Corrinoid-binding protein; MtxA: Methylcorrinoid:carbon-carrier methyltransferase; RamX: reductive activase/activating enzyme for corrinoid-binding protein MtxC; geTMM: Gene length corrected trimmed mean of M values; GTDB: Genome taxonomy database

## Acknowledgments

This research is a contribution of the EMERGE Biology Integration Institute funded by the NSF Biology Integration Institutes Program, Award #2022070. JBE was fully, and MAB, KCW, DRC, VF-Z, RKV, JPC, BJW, MMT, and VIR were partially supported by this award. We thank the Swedish Polar Research Secretariat and SITES for the support of the work done as Abisko Scientific Research Station. SITES is supported by the Swedish Research Council’s grant 4.3-2021-00164. This study was partially funded by the Genomic Science Program of the United States Department of Energy Office of Biological and Environmental Research, grant #DE-SC0010580. BBM and KCW are also supported by an Early Career Award to KCW from the National Science Foundation (NSF) under award number 1912915. A portion of this research was performed under the Facilities Integrating Collaborations for User Science (FICUS) initiative and used resources at the DOE Joint Genome Institute and the Environmental Molecular Sciences Laboratory, which are DOE Office of Science User Facilities. Both facilities are sponsored by the Office of Biological and Environmental Research and operated under Contract Nos. DE-AC02-05CH11231 (JGI) and DE-AC05-76RL01830 (EMSL). This includes EMSL projects (10.46936/sarr.proj.2018.50229/60000028 and 10.46936/lser.proj.2021.51858/60000347) awarded to PI KCW. Autochamber measurements between 2013 and 2017 were supported by a grant from the US National Science Foundation MacroSystems program (NSF EF 1241037, PI Varner). LC MS/MS analysis was conducted at the Analytical and Biological Mass Spectrometry (ABMS) Core Facility personnel at the University of Arizona supported by the RII (Research, Innovation, and Impact) and TRIF (Technology and Research Initiative Fund) initiative. A portion of the MAG recovery was performed on the Ohio Supercomputer Center (119)

We thank Adrienne Narrowe and Emily Bechtold for methods/analysis consultations, and Tyson Claffey and Richard Wolfe for Colorado State University server management.

## Author’s contributions

*Conceptualization: JBE, MAB, KCW*

*Methodology: JBE, KCW, MAB, BBM, DWH, VFZ, MMT, DRC, VIR, JPC, CKM, BJW, GWT, RKV, PMC, RAW*

*Software: MAB Validation: MAB*

*Formal analysis: JBE, KCW, MAB, BBM, DWH, VFZ, DRC, JPC, CKM, BJW, RKV, PMC, RAW, MMT*

*Investigation: JBE, KCW, BBM, DWH, VFZ, DRC, JPC, CKM, RKV, PMC, RAW, BJW*

*Resources: KCW, DWH, MMT, VIR, JPC, CKM, GWT, Rkv*

*Data Curation: JBE, KCW, MAB, BBM*

*Writing-Original Draft: JBE, MAB, KCW, BBM*

*Writing - Review and Editing: All authors read and provided feedback/edits on the draft manuscript and approved its final form*.

*Visualization: JBE*

*Supervision: JBE, KCW, MMT, VIR, RKV, JPC, CKM, GWT*

*Project administration: JBE, KCW, MMT, VIR, RKV, JPC, CKM, GWT, BJW, PMC, RAW Funding acquisition: KCW, MMT, VIR, JPC, BJW, GWT, RKV*

## Supplementary Figures and Legends

**Supplemental Figure 1.**
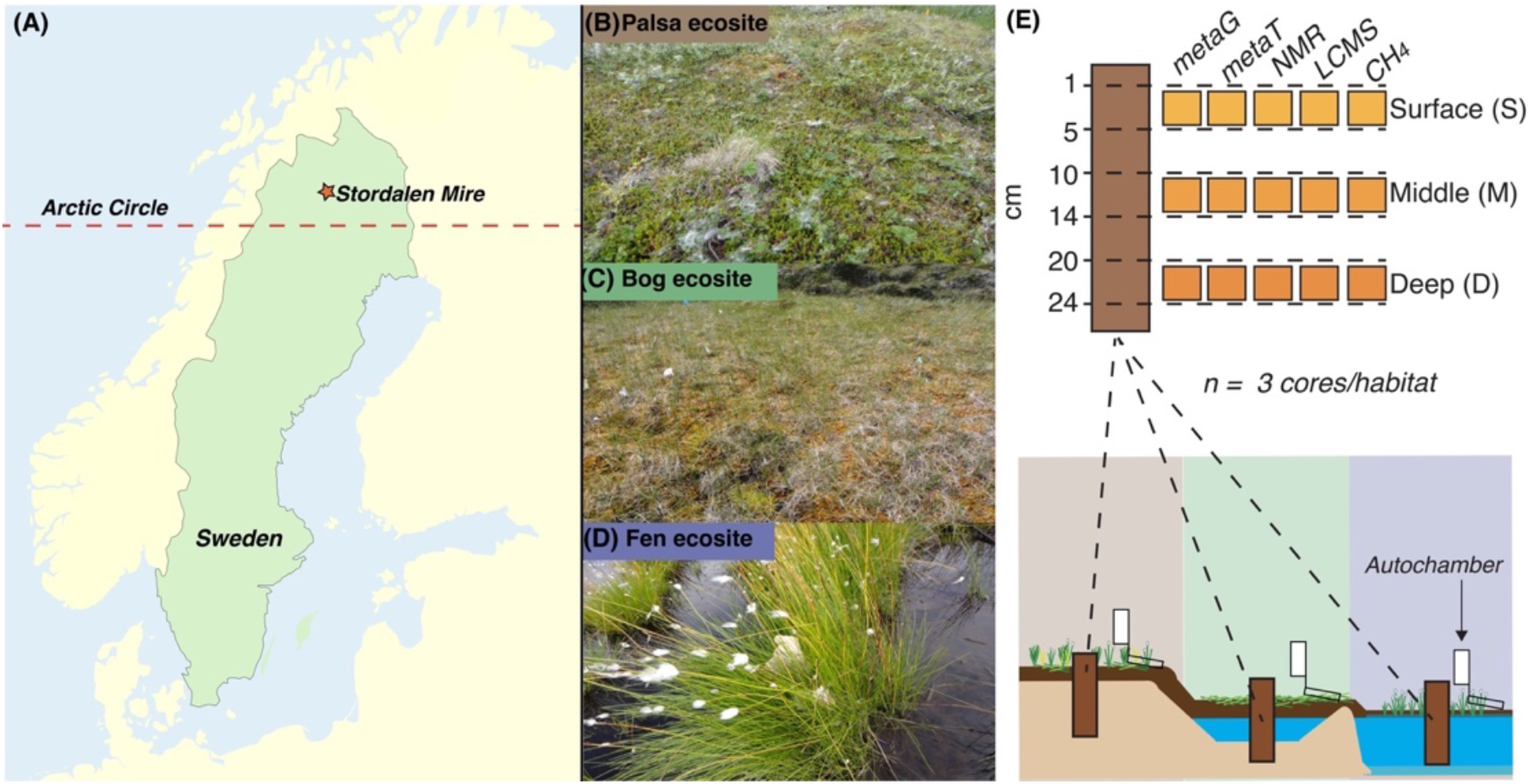
Stordalen Mire is a thawing permafrost peatland in Arctic Sweden. (A) Map showing the location of Stordalen Mire within the Arctic Circle in Northern Sweden. The Mire contains three distinct habitat types which constitute a discontinuous natural permafrost thaw gradient, across which the seasonally thawed active layer was sampled: (B) Palsa, representing frozen intact permafrost, (C) Bog, often formed from partially thawed permafrost with a perched water table and (D) Fen, often formed from fully thawed permafrost with a saturating water table. (E) Sampling design for July 2016; triplicate soil cores were obtained from each habitat and subdivided into three depths (surface = 1-5 cm, middle = 10-14 cm, deep = 20-24 cm) for analyses including metagenomics, metatranscriptomics, NMR, LCMS. Both methane flux data and porewater methane measurements were obtained from or near coring sites. All photographs (B-D) were taken by Nicole Raab in July of 2016, and retrieved from the EMERGE database.

**Supplemental Figure 2.**
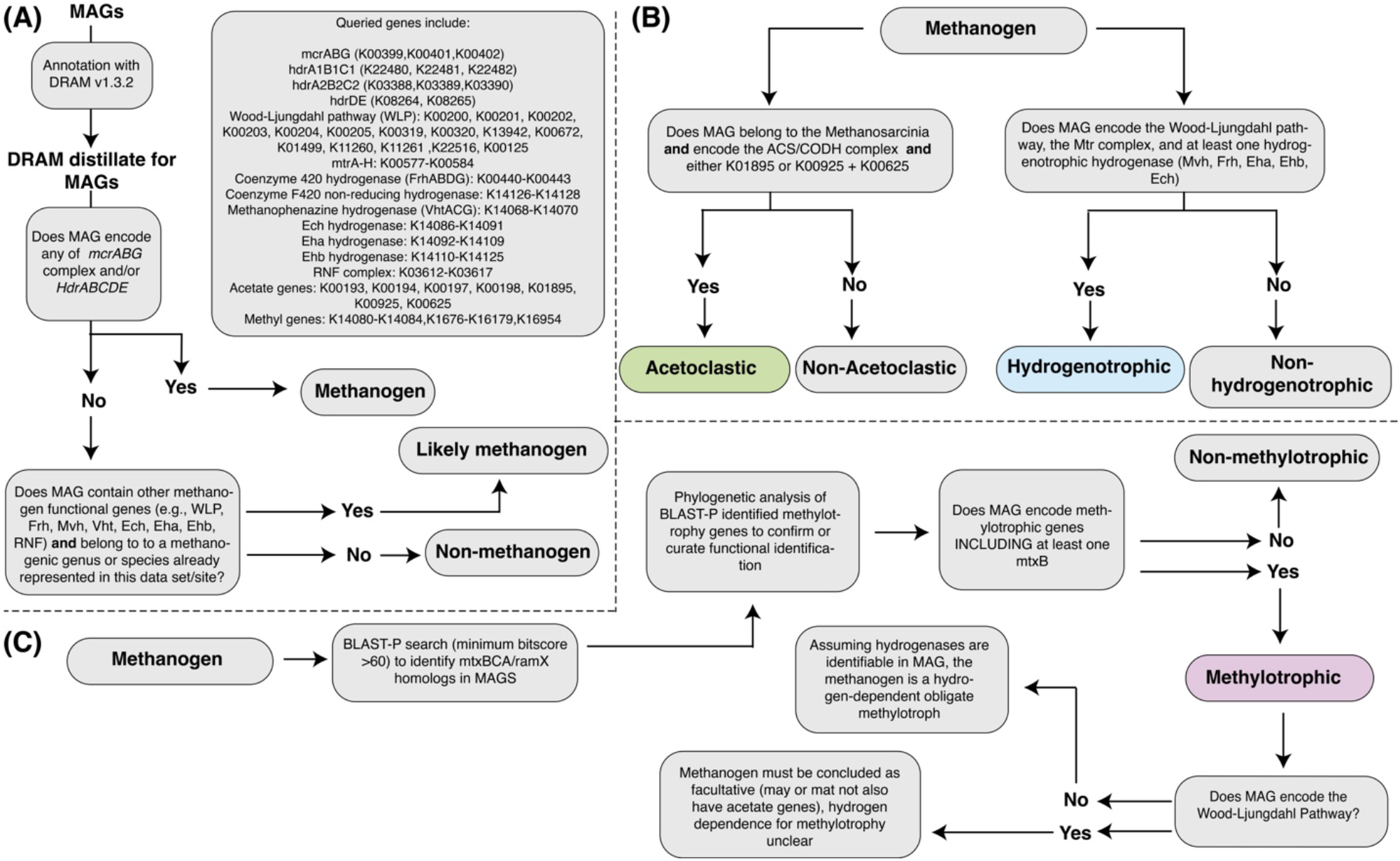
Workflow for curation of methanogen physiology. (A) Method for identifying MAGs as encoding potential for methanogenesis per analysis of DRAM annotation/distillate (B) Method used for identifying MAGs as encoding potential for either acetoclastic or hydrogenotrophic methanogenesis per DRAM output (C) Workflow used to curate methylotrophic potential across methanogen MAGs independent of DRAM/bioinformatic databases. Note that B and C are not mutually exclusive, and some MAGs might meet the criteria for encoding multiple pathways of methanogenesis.

**Supplemental Figure 3.**
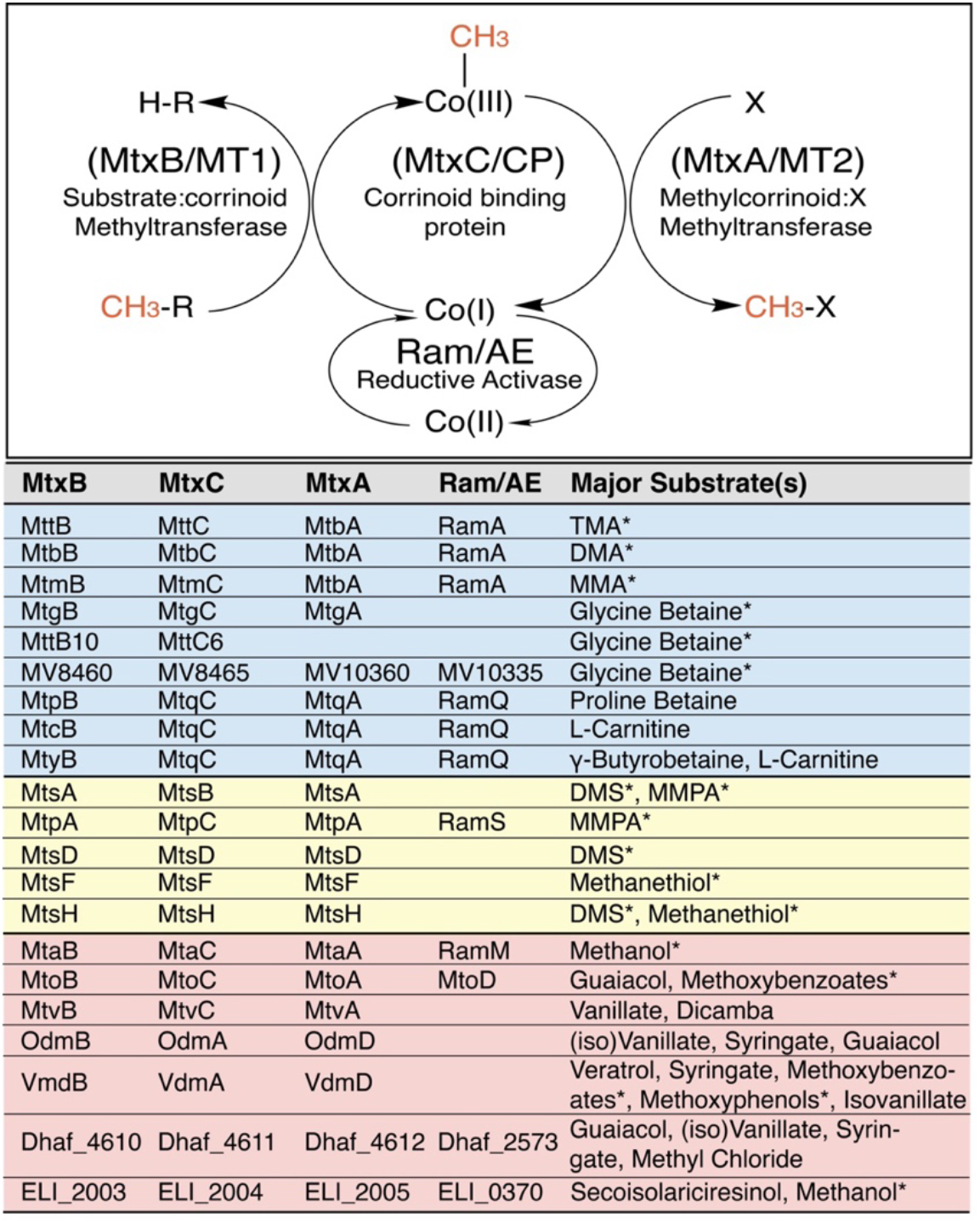
List of all biochemically verified corrinoid dependent methyltransferase systems. List of substrate-specific protein names/identifiers used in relevant literature for all well characterized corrinoid-dependent methyltransferase systems. Major substrates (methyl-N = blue, methyl-S = yellow, methyl-O = red) are provided for each set of proteins, and these represent those found to be kinetically comparable substrates for methyltransferase that would be reasonable to suspect as *in vivo* substrates. This list includes proteins described from both methanogens and acetogens; substrates with an asterisk next to them have been specifically shown to support direct methanogenesis, versus bacterial methylotrophy/acetogenesis.

**Supplemental Figure 4.**
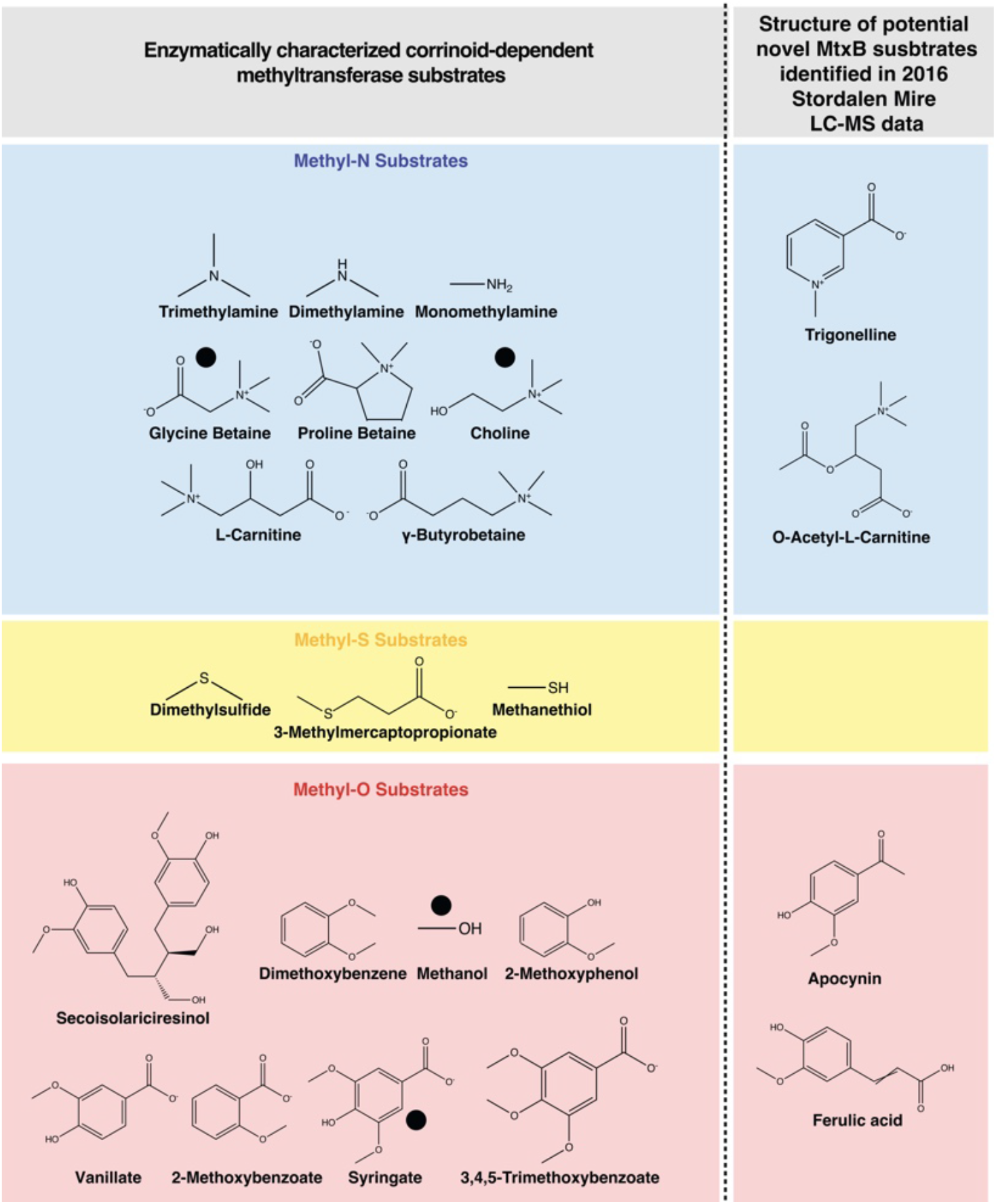
Structures of known and proposed methylotrophic substrates found in Stordalen Mire. To the left of the dashed line are all so far biochemically validated substrates for corrinoid-dependent methyltransferase systems utilized by methylotrophic methanogens, acetogens, and other bacteria. All have been enzymatically profiled except choline, which has so far only been shown at the culture-level to support direct methanogenesis via presumed methylotrophy. Some of these substrates – denoted with a black dot - were found present in Stordalen Mire via metabolite analyses. Note that *D*- but not *L*-Carnitine was identified via LC-MS, therefore the compound is not marked with a dot. Shown to the right of the dashed line are the structures of all methylated compounds identified via LC-MS in the Mire which have not been previously shown to, but are here purported to, support methylotrophy based on structural similarity to known substrates.

**Supplemental Figure 5.**
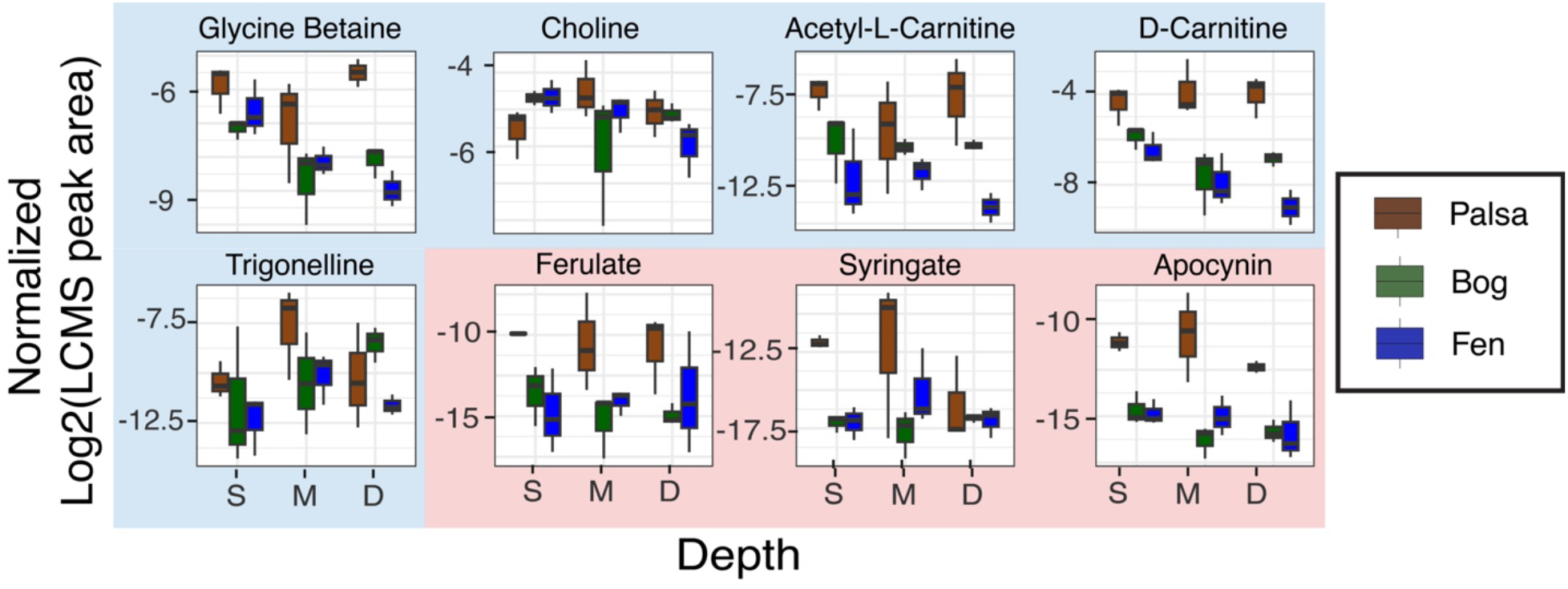
Individual LC-MS peak area plotted for the methylated compoundsa across habitats. Boxplots show the normalized individual LC-MS peak areas within each habitat and depth for all methylated compounds identified from Stordalen Mire. The categorically (methyl-N, methyl-O) summed averaged peak areas of these are plotted in Figure 3B.

**Supplemental Figure 6.**
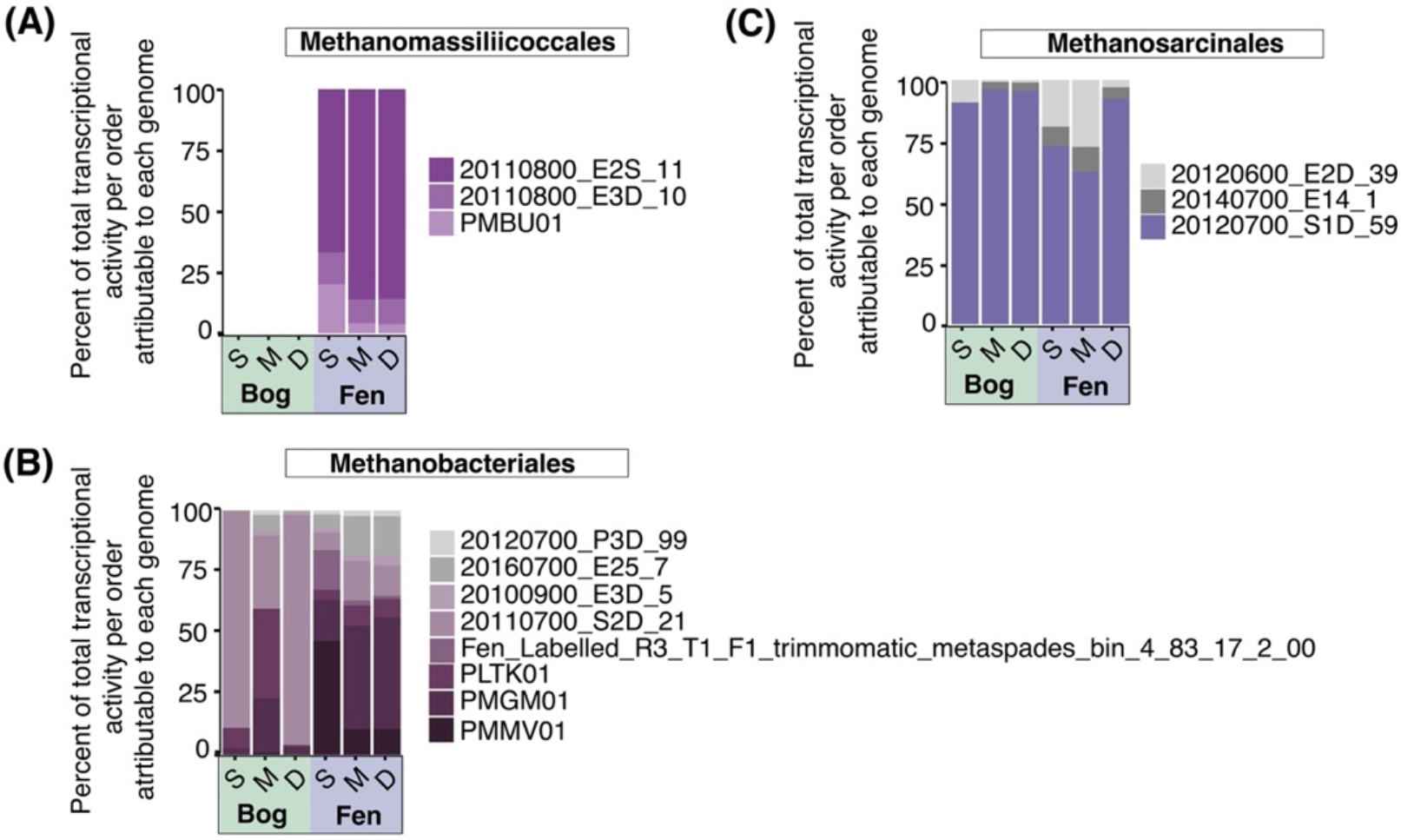
Actively methylotrophic methanogens consistute most of the activity of their respective orders. Stacked bar chart showing the relative contribution of individual active MAGs to the summed transcriptional activity of each methylotrophic order across habitat and depth. Bars are colored purple if MAGs met a threshold of being considered active methylotrophs, meaning that they were found to express at least a substrate-demethylating methyltransferase (*mtxB*) and either an *mtxC* or *mtxA* homolog. Bars representing MAGs that did not meet this threshold are colored in grey.

**Supplemental Figure 7.**
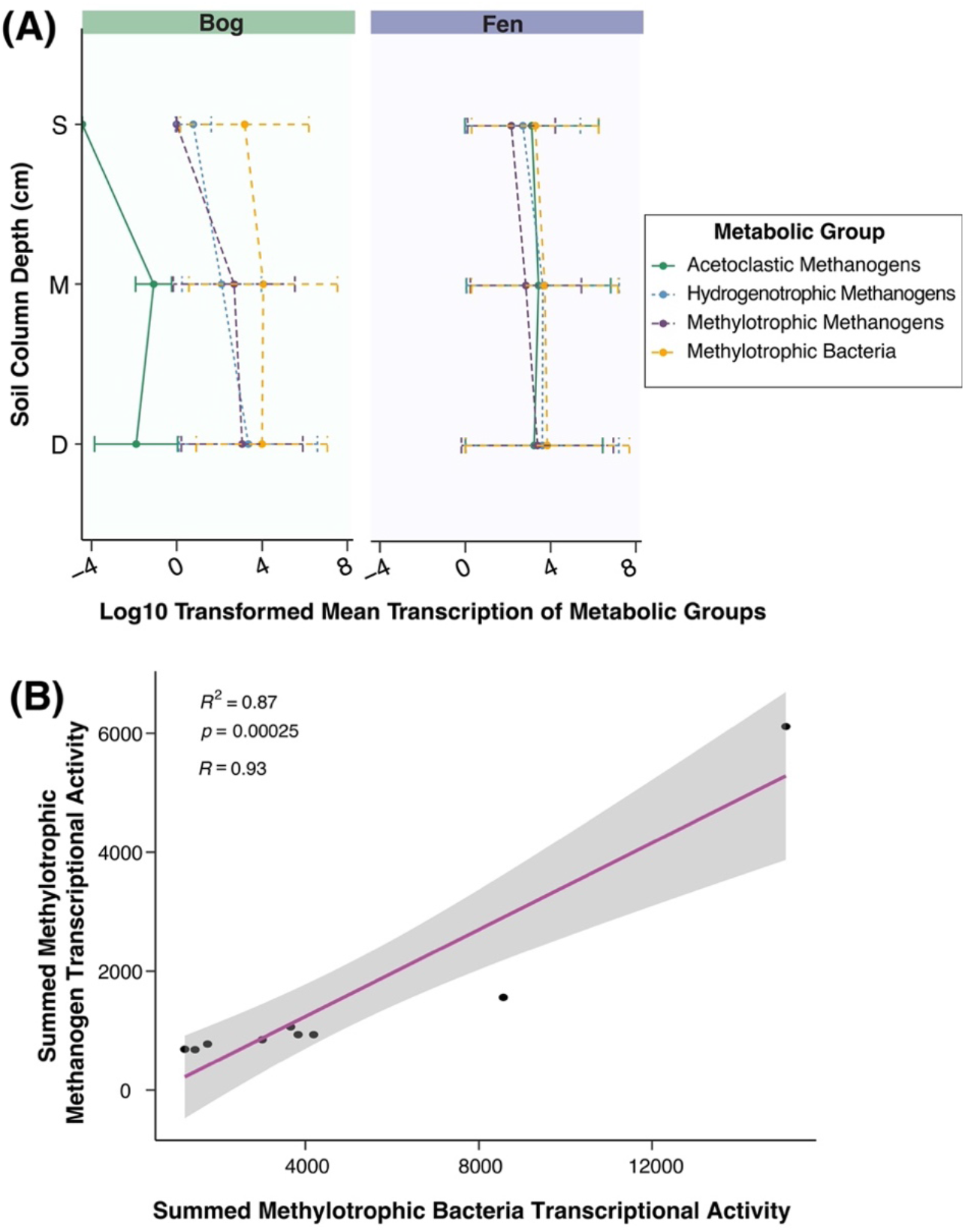
Co-occurrence of methanogen and methylotrophic bacteria activity. (A) Line plots showing the average summed transcriptional activity of each methylotrophic bacteria, methylotrophic methanogens, acetoclastic methanogens, and hydrogenotrophic methanogens in the bog and the fen. Error bars represent one standard deviation. Acetoclastic methanogens appear to be less active in the shallower bog, but generally the four metabolic groups are simultaneously active in the same habitats and depths.(B) Within the fen across depths, a strong positive correlation was seen between the mean summed transcriptional activity of methylotrophic methanogens and methylotrophic bacteria.

## Supplemental Table Legends

**Table S1.** Methanogen metagenome assembled genome quality data, and accession information for metagenome and metatranscriptome data sets.

**Table S2.** Methanogen metagenome assembled genome physiological curation data

**Table S3.** Literature based knowledge/references on corrinoid-dependent methyltransferase system genes, plus FASTA sequences used in phylogenetic analyses.

**Table S4.** Metabolite data used in paper including field methane measurements, plus subsets of NMR and LCMS data.

**Table S5.** Table with relative abundance calculations for methanogen genomes in metagenome and metatranscriptome data sets.

**Table S6.** Data supporting physiological curation of bacterial methylotrophs.

